# Hundreds of antimicrobial peptides create a selective barrier for insect gut symbionts

**DOI:** 10.1101/2023.10.16.562546

**Authors:** Joy Lachat, Gaëlle Lextrait, Romain Jouan, Amira Boukherissa, Aya Yokota, Seonghan Jang, Kota Ishigami, Ryo Futahashi, Raynald Cossard, Delphine Naquin, Vlad Costache, Luis Augusto, Pierre Tissières, Emanuele G. Biondi, Benoît Alunni, Tatiana Timchenko, Tsubasa Ohbayashi, Yoshitomo Kikuchi, Peter Mergaert

## Abstract

The spatial organization of gut microbiota is crucial for the functioning of the gut ecosystem, although the mechanisms that organize gut bacterial communities in microhabitats are only partially understood. The gut of the insect *Riptortus pedestris* has a characteristic microbiota biogeography with a multispecies community in the anterior midgut and a mono-specific bacterial population in the posterior midgut. We show that the posterior midgut region produces massively hundreds of specific antimicrobial peptides (AMPs), the Crypt-specific Cysteine-Rich peptides (CCRs) that have membrane-damaging antimicrobial activity against diverse bacteria but posterior midgut symbionts have elevated resistance. We determined by transposon-sequencing the genetic repertoire in the symbiont *Caballeronia insecticola* to manage CCR stress, identifying different independent pathways, including novel AMP-resistance pathways unrelated to known membrane homeostasis functions as well as cell envelope functions. Mutants in the corresponding genes have reduced capacity to colonize the posterior midgut, demonstrating that CCRs create a selective barrier and resistance is crucial in gut symbionts. Moreover, once established in the gut, the bacteria differentiate into a CCR-sensitive state, suggesting a second function of the CCR peptide arsenal in protecting the gut epithelia or mediating metabolic exchanges between the host and the gut symbionts. Our study highlights the evolution of an extreme diverse AMP family that contributes to establish and control the gut microbiota.

## Introduction

The animal gut is colonized by bacterial communities, which provide essential functions to the host (1, 2). The phylotype richness and total abundance of this gut microbiota varies strongly among the animals form low to extraordinarily high (1). Moreover, in animals ranging from humans to insects, gut microbiota do not constitute a homogeneous mixture but are spatia lly organized and form discrete bacterial communities located in specific microhabitats along the longitudinal and transverse axes of the gut (3–6). How this microbial biogeography is established is only partially understood but is potentially correlated with physical barriers such as mucus, peritrophic membrane and crypts, gradients of chemical parameters such as pH or oxygen levels, bacteriophages and nutrient availability as well as host immune effectors. Among the latter are antimicrobial peptides (AMPs), which are secreted in the gut lumen and come in contact with the microbiota (7–10). AMPs contribute to establish an epithelia - microbiota equilibrium along the transverse axis of the gut by regulating the species composition and location of the microbiota according to the resistance and sensitivity patterns of its members (10). Thus, gut commensals are expected to be resilient to AMPs (11, 12) but how they adapt and how important this adaptation is for colonization of their specific niche within the gut remains largely unexplored. Moreover, it is not known if AMPs exert control on the spatial organization of microbiota along the longitudinal gut axis.

The bean bug *Riptortus pedestris* has a particular midgut organization, associated with a simple microbiota displaying a characteristic biogeography. The midgut has four morphologically and functionally distinct compartments, labelled M1 to M4. The anterior M1 to M3 regions are involved in food digestion and have a variable and transient microbiota, which is ingested through feeding. The posterior M4 region on the other hand, composed of two rows of crypts branched on a central tract, does not contribute to food digestion and is associated with a stable, (nearly) mono-specific and high-abundant microbiota that is also acquired from the environment and sorted out from the M3 microbiota (13). The M4 bacteria are very specific, belonging to the *Caballeronia* genus and are mostly present as a single colonizing species, established through a multifaceted selection process. A sorting organ located at the entry of the M4 region winnows out the M3 microbiota allowing only a subset of species to enter the M4 (14). After a successful initial passage of bacteria through the soring organ and infection of the M4 region, secondary infections are inhibited by closure of the sorting organ (15). The infecting bacteria induce in the M4 crypts developmental processes, including oxygenation by tracheal formation (16) and the maturation of the crypts by intestinal stem cell stimulation and apoptosis inhibition that creates the luminal space in the crypts for bacterial colonization (17). Finally, microbe-microbe competition within the crypts results in the elimination of the least adapted strains and the dominance of a single strain in the M4 region (18). Among the bean bug colonizers, *Caballeronia insecticola* has emerged as a model species (19). We took advantage of this simplified gut-microbe interaction model to explore if together with the already known mechanisms, AMP challenge contributes to create the gut biogeography in *R. pedestris* and if AMP resistance in *C. insecticola* is crucial for M4 crypt colonization.

## Results

### The *Riptortus pedestris* midgut expresses hundreds of AMP-like genes

A preliminary transcriptome analysis of the M4 midgut region has identified a novel class of secretory peptides, which we call the CCRs (20, 21). In order to define the expression pattern of *CCR* genes, the transcriptome was determined by RNA-seq in midgut regions of insects that were reared for different times in the presence or absence of the *C. insecticola* gut symbiont (Fig. 1A). The pooled sequencing reads were assembled in a set of unique transcripts and encoded proteins. Hidden Markov Models based on the previously identified CCR sequences were used to identify in the newly generated transcriptome the complete set of CCR sequences. This analysis revealed 310 *CCR* transcripts (SI Appendix, Data S1). The encoded CCR peptides do not show high similarity apart from a pattern of conserved cysteine residues (Fig. 1B). Despite their sequence divergence, AlphaFold2 predicted similar folds for tested CCR peptides, consisting of three pairs of -sheets that are probably connected by cystine bridges (Fig. 1C). Differential expression analysis revealed that the majority of the *CCR* genes are most strongly expressed in the midguts of symbiotic insects (Fig. 1D and SI Appendix, Data S1). Subsets of genes were specific for the M3, M4B and the majority for the M4 region carrying the *C. insecticola* bacteria, suggesting that the encoded peptides target the symbionts. Moreover, the *CCRs* are among the most strongly expressed transcripts in the overall transcriptome (Fig. 1E), suggesting a primordial role of the peptides in the midgut. The *CCR* genes did not exhibit similarity to known sequences of other organisms. However the taxonomica l ly restricted nature of the genes as well as the structure of the CCRs, being small, secreted and characterized by conserved cysteine residues, remind strongly to AMP gene families (10) and AMP prediction tools confirmed this presumption (Fig. 1C and SI Appendix, Data S1). Whole mount *in situ* hybridization with the infected-M4-specific gene *CCR0043* showed that the gene is expressed uniformly by the epithelial cells in all M4 crypts (Fig. 1F and SI Appendix, Fig. S1). This pattern contrasts with the mammalian small intestine where specialized cell types at the base of crypts express AMP genes (22).

**Figure 1.**
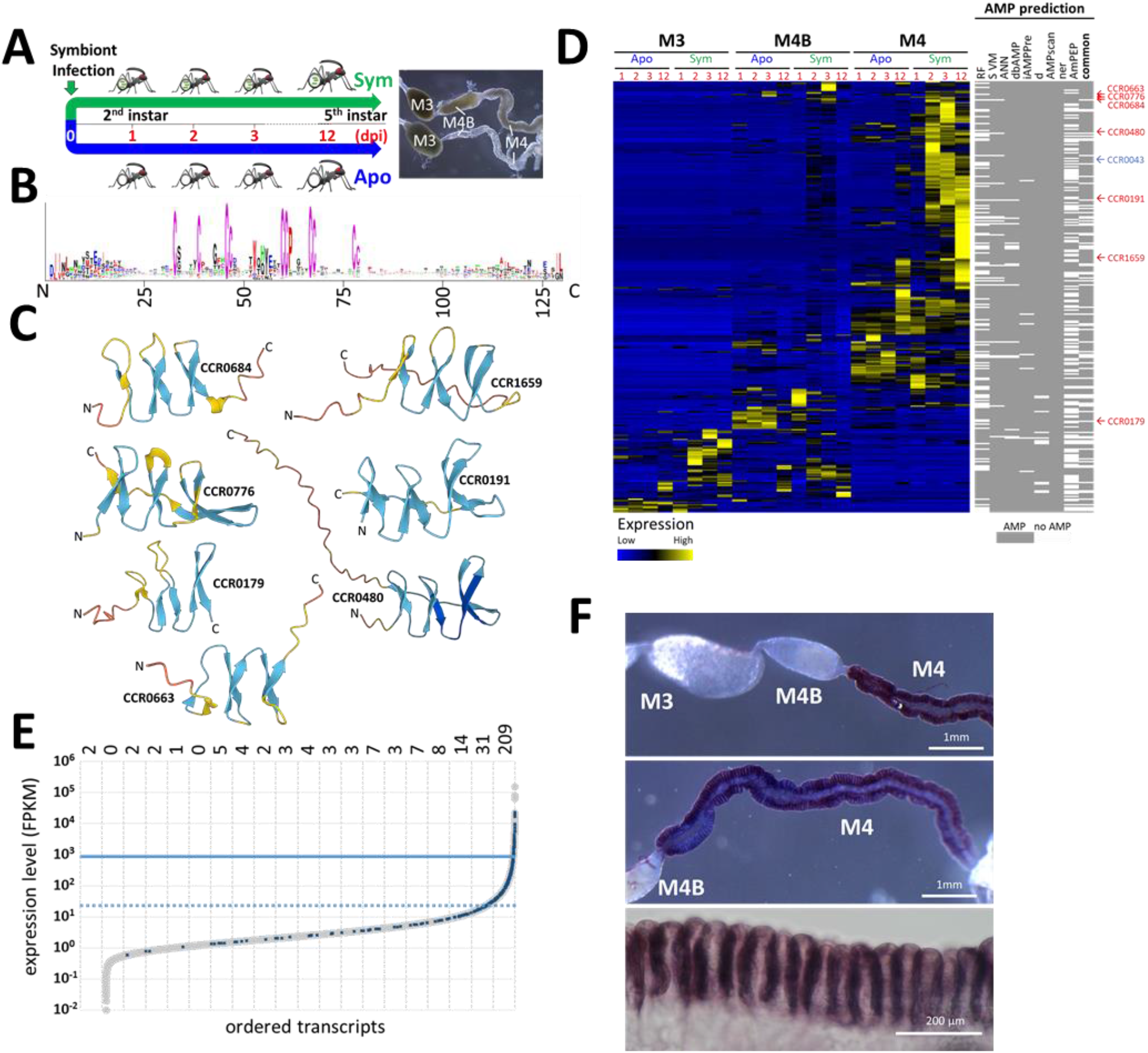
CCRs are symbiosis-specific AMP-like peptides. (**A)** Experimental setup for transcriptome analysis. First day second instars were divided in two groups. To one of them, *C. insecticola* symbionts were administered (green, Sym) and the other group remained free of symbionts (blue, Apo). Insects were dissected in the second (1, 2, and 3 days post inocula t ion [dpi]) or fifth instar (12 dpi) and the M3, M4B and M4 regions were harvested for transcripto me analysis. The pictures at the right show representative guts of a Sym insect at 3 dpi (top) and a same age Apo insect (bottom). (**B**) Logo profile of the mature CCR peptides identified in the transcriptome, highlighting the sequence diversity of the peptides and the ten conserved cysteine residues. (**C**) AlphaFold2 structural predictions of examples of CCR peptides showing antiparallel -sheets carrying the cysteine residues. (**D**) Blue-black-yellow heat map of the relative expression profile of the identified CCR genes and white-grey heat map of AMP predictions. Sample identity in the expression heat map is indicated at the top and is according to panel A. AMP prediction tools are Random Forest (RF), Support Vector Machine (SVM), Artificial Neural Network (ANN), AMPpredictor (dbAMP); iAMPpred; Antimicrobial Peptide Scanner (AMPscanner); (AmPEP_v1 and AmPEP_v2). A consensus prediction (6 out of 7 positive predictions) is indicated in the last column. The peptides used for functio nal characterization are indicated at the right of the heat maps. (**E**) Whole-mount *in situ* hybridization with a *CCR0043* antisense probe on the dissected midgut of a 3 dpi symbiot ic insect. Positive signal appears with a blue-brownish color. CCR0043 is specifically expressed in the M4 and uniformly in all crypts. Control *in situ* hybridizations on the gut of aposymbiot ic insects and with a sense probe on symbiotic insects are shown in Supplemental Figure S2. (**F**) *CCR* transcript expression levels. Transcripts are ordered according to their expression level in the x-axis and their expression levels (FPKM, Fragments Per Kilobase of transcript per Millio n mapped reads) are plotted in the y-axis. All transcripts are indicated with grey dots and the *CCR* transcripts are indicated with blue crosses. The dotted and plain blue horizontal lines correspond to the mean expression level of all transcripts and *CCR* transcripts, respectively. The numbers above the plot indicate the number of *CCR* transcripts present in 5-percentile bins of transcripts. 77 % of the *CCR* transcripts are among the 10 % highest expressed transcripts in the midgut.

### CCRs have antibacterial activity but gut colonizers are resistant

Based on the expression pattern and the predicted AMP activity, we selected CCRs for chemical synthesis (Fig. 1C and SI Appendix, Table S1). These seven CCRs, together with thanatin and riptocin, two known innate immunity-related AMPs of *R. pedestris* (23), LL37 and NCR335, from mammal and plant origin respectively (24, 25) and bacterial polymyxin B (PMB), were tested for growth inhibiting activity against a panel of taxonomically diverse bacterial species *Bacillus subtillis*, *Sinorhizobium meliloti*, *Paraburkholderia fungorum* and *C. insecticola*. The first two species are unable to colonize the *R. pedestris* midgut while the latter two can efficiently proliferate in the crypts (18). In agreement with the bioinformatics predictions, the CCRs had growth inhibiting activity against *B. subtilis* and *S. meliloti* although with variable strengths (Fig. 2A, B). On the other hand, the two species, *P. fungorum* and *C. insecticola*, that are able to colonize the gut crypts, are not or only weakly affected by the tested CCRs (Fig. 2A, B). This pattern of sensitivity/resistance to CCRs matches with the response of these species to PMB and in part to the other tested peptides.

**Figure 2.**
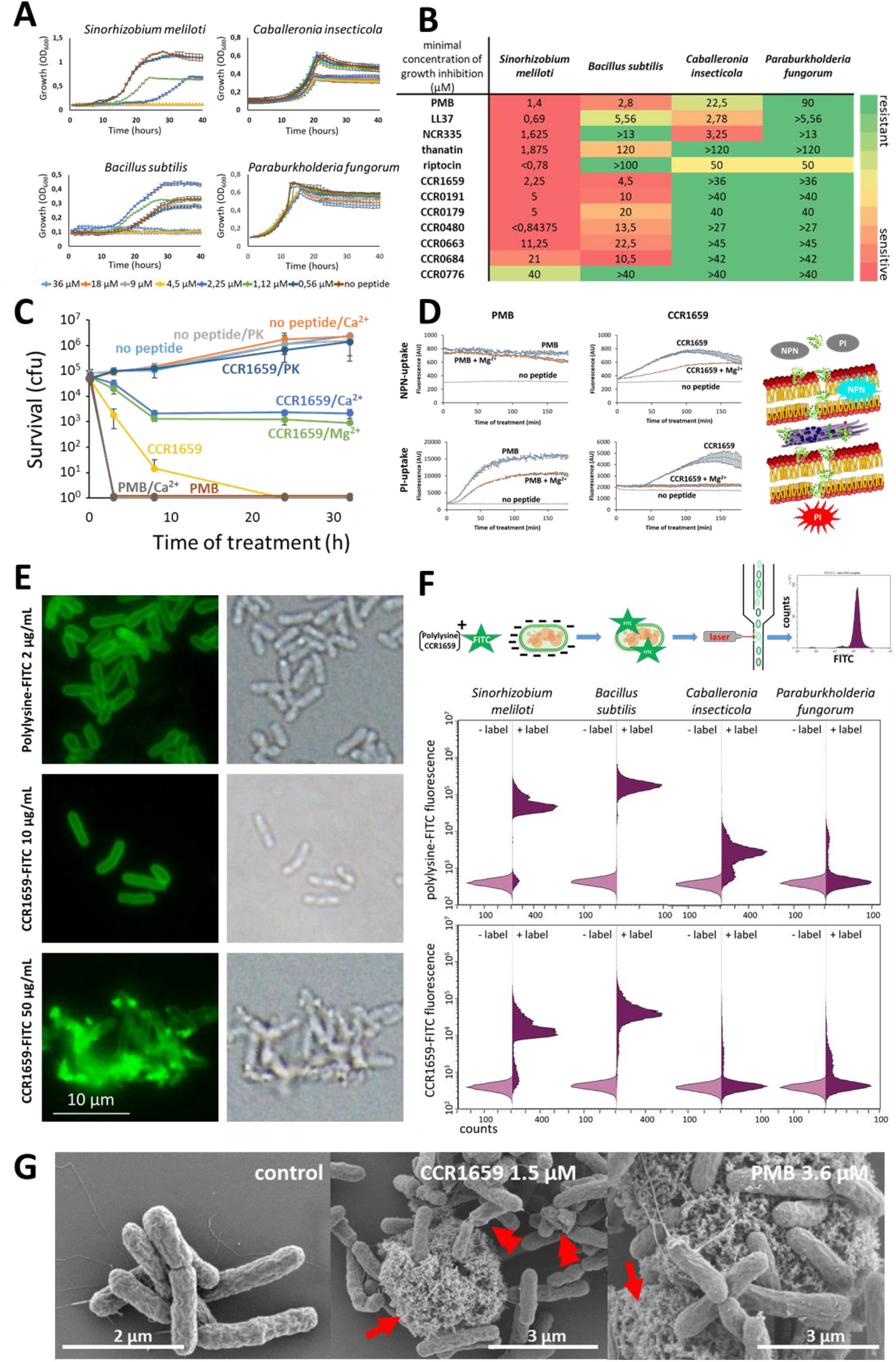
CCR peptides are membrane-targeting AMPs. (**A**) Growth inhibition of the indicated bacterial species by different concentrations of CCR1659. Error bars are standard deviations (n=3). (**B**) Minimal concentrations (in µM) of growth inhibition of the indicated strains by various peptides. (**C**) Bactericidal activity of 25 µM CCR1659 and 25 µM PMB. PK: proteinase K; Ca^2+^: activity in the presence of 5 mM CaCl_2_; Mg^2+^ activity in the presence of 5 mM MgCl_2_. Error bars are standard deviations (n=3). (**D**) NPN and PI uptake by *S. meliloti* cells in response to treatment with 10 µM CCR1659 or 10 µM PMB in the presence or absence of 5 mM MgCl_2_. NPN is a lipophilic dye that fluoresces in hydrophobic environments such as bacterial phospholipids exposed by outer membrane damage; PI is a membrane impermea nt DNA-intercalating dye that fluoresces upon DNA binding in the cytoplasm, indicative of permeabilisation of both the outer and inner membrane. Error bars are standard deviatio ns (n=3). (**E**) Fluorescence microscopy (left) of *S. meliloti* cells treated with Polylysine-FITC or CCR1659-FITC at the indicated concentrations. Corresponding bright field images are shown at the right. (**F**) Flow cytometry analysis of Polylysine-FITC (top) or CCR1659-FITC binding by the indicated strains. Light purple histograms are control measurements without fluoresce nt label (-label); the dark purple histograms are in the presence of the fluorescent label (+label). (**G**) SEM micrographs of untreated *S. meliloti* cells (left) or treated with 1.5 µM CCR1659 (middle) or with 3.6 µM PMB (right). The arrows indicate cellular material released from cells. The double arrowheads indicate cells with lost turgor.

### CCRs have membrane-damaging bactericidal activity

CFU counting showed the bacterial reduction from 10^7^ CFU to no colonies after treatment of sensitive *S. meliloti* with the CCR1659 peptide for a few hours, indicating that the growth inhibition results from a bactericidal activity, similarly as for PMB (Fig. 2C). The bactericidal activity of CCR1659 was abolished by prior Proteinase K treatment of the peptide and inhibited by the presence of the divalent cations Ca^2+^ and Mg^2+^, which interfere with the electrostatic interaction of AMPs with negatively charged membrane lipids and diminish the activity of membrane-targeting AMPs (26) (Fig. 2C; SI Appendix, Fig. S2). To acquire insight in the killing mode of CCR1659, we tested the hypothesis that the peptide disrupt bacterial membranes, like PMB and the other tested AMPs do (27–29). Outer and inner membrane integrities in *S. meliloti* were consecutively damaged by both CCR1659 and PMB treatment, as measured respectively by 1-N-PhenylNaphthylamine (NPN) and Propidium Iodide (PI) uptake leading to enhanced fluorescence (Fig. 2D). In agreement with the membrane disruption, fluorescence microscopy showed that FITC-modified CCR1659 labelled the envelope of *S. meliloti* cells in a similar way as polylysine-FITC, which is a polycation known to interact with negatively charged membranes of bacteria (30, 31) (Fig. 2E). Binding of CCR1659 to the envelope suggests that its killing efficiency depends on the strength of envelope binding. To test this assumption, we measured with flow cytometry the binding level of CCR1659-FITC and polylysine-FITC to the above panel of species. Strikingly, CCR1659-sensitive *S. meliloti* and *B. subtilis* were strongly labeled with these two molecules while resistant *C. insecticola* and *P. fungorum* only weakly (Fig. 2F). Thus, the level of binding to cells is correlated with the susceptibility/resistance pattern. Scanning electron microscopy (SEM) of CCR1659-treated *S. meliloti* cells further confirmed the membrane-perturbing activity of the peptide that provoked the leakage of fibrous materials from damaged cells similarly as PMB (Fig. 2G; SI Appendix, Fig. S3). Together, this data reveal that the M4 symbiotic region of the gut produces a remarkably large arsenal of CCR peptides with membrane-damaging AMP activity.

### The *Caballeronia insecticola* genetic repertoire determining AMP resistance

Species that colonize the midgut display a high level of resistance to CCRs and other AMPs suggesting that resistance is a prerequisite for efficient gut colonization. To test this hypothesis, we aimed to identify the resistance determinants in *C. insecticola* and assess if they control gut colonization. A transposon mutant library (32) was used to perform a Tn-seq screen with PMB, since PMB has a similar membrane action as CCR peptides and is commercially accessible in sufficient quantities for Tn-seq experiments. The screen, performed with three sub-lethal PMB concentrations, resulted in 54 genes whose mutation provoked a fitness defect with the highest concentration. With the lower PMB concentrations, subsets of these genes were identified suggesting a multifactorial resistance with some mechanisms contributing more strongly than others (Fig. 3A, B; SI Appendix, Fig. S4 and Data S2). In agreement with the membrane - targeting mode of action of PMB, the majority of fitness genes are involved in the generation of bacterial envelope components, including LPS, peptidoglycan, phospholipids, hopanoids and membrane protein machineries. In order to validate the Tn-seq results, we constructed insertion and deletion mutants in 11 genes selected among the 54 PMB fitness genes. These genes are predicted to be involved in the biosynthesis of the LPS core (*dedA*, *waaC* and *waaF*) (33–35), LPS O-antigen (*wbiF*, *wbiG*, *wbiI*, *wzm* and *rfbA*) (36), peptidoglycan (*dedA*), membrane protein machineries (*tolB* and *tolQ*) (37, 38), in addition to a gene (*tpr*) encoding a tetratricopeptide repeat protein of unknown function. Complementing strains were constructed for some of the mutants. Sensitivity assays with PMB and colistin (COL), another polymyxi n-family AMP, confirmed that each mutant had an 8- to 32-fold increased sensitivity compared to the WT (Fig. 3C) while the complemented mutants were restored to WT-levels (SI Appendix, Fig. S5). Thus, the Tn-seq analysis correctly identified genetic determinants for PMB resistance in *C. insecticola*.

**Figure 3.**
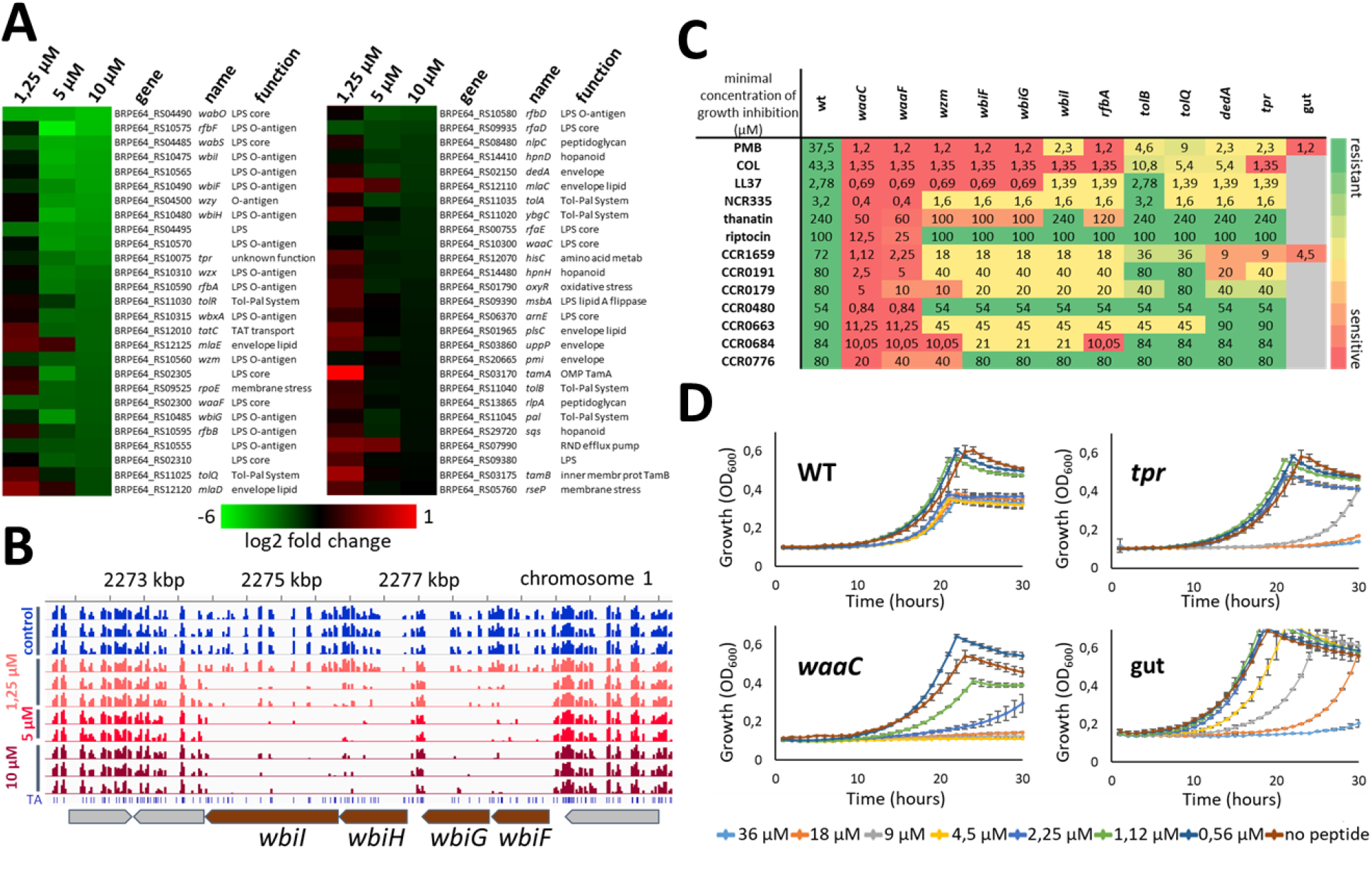
Identification of AMP resistance genes by Tn-seq. (**A**) Heat map showing the level of depletion of transposon insertions in the indicated genes in the *C. insecticola* population grown in the presence of PMB at the indicated concentrations. The color-code scale indicates the log2 fold change in the insertion abundance under the test conditions relative to the control conditions. (**B**) IGV view of Tn-seq sequencing data for a selected genomic region of *C. insecticola*. The histograms indicate the abundance of mutants in the Tn-seq population for the indicated samples. Genes whose products contribute to PMB resistance have a lower frequency of Tn insertions in peptide treatment screens than in the control. (**C**) Mutants in selected genes are hypersensitive to AMPs. Heat map and minimal concentrations of growth inhibition of the indicated wild-type and mutant strains by the listed peptides. Minimal concentrations are indicated in µM. The color key of the heat map is as indicated at the right. Grey cells indicate not tested. (**D**) Growth inhibition of the indicated strains by different concentrations of CCR1659. Gut indicates crypt-colonizing *C. insecticola* bacteria, directly isolated from dissected M4. Error bars are standard deviations (n=3).

In line with the sensitivity of the mutants to PMB and the membrane-attacking properties of CCRs, we found that all mutants were more sensitive than WT for at least one of the tested CCR peptides and the other available AMPs (Fig. 3C, D). The *tolB* and *tolQ* mutants were the least sensitive and displayed only a slight difference compared to WT for all tested peptides. The *dedA* and *tpr* mutants were strongly affected by the CCR1659 peptide (Fig. 3D) and moderately by the other tested peptides. The mutants *wzm*, *wbiF*, *wbiG*, *wbiI* and *rfbA* were sensitive to several of the tested CCRs although in many cases, enhanced sensitivity was not resulting in a complete growth inhibition but in a retarded and lesser growth compared to untreated control and the WT grown with the same peptide concentration. The *waaC* and *waaF* mutants were the most strongly affected, being more sensitive than WT to all tested peptides and at higher peptide concentrations, their growth was completely blocked (Fig. 3D). Taken together, the *C. insecticola* genes that were revealed by the PMB Tn-seq screen, contribute also to resistance towards other membrane-attacking AMPs, including the CCRs.

### Different pathways contribute to AMP resistance in *Caballeronia insecticola*

Because the tested AMPs interfere with bacterial membrane function, we characterized the cell envelope of the mutants. Since some of the mutated genes are known or suspected to be involved in LPS biosynthesis, we analyzed the LPS structure of all mutants by PAGE profiling and by mass spectrometry analysis of their lipid A moiety, which is proposed to be a direct target of PMB (27, 28) (Fig. 4A, B). The *tpr*, *dedA* and *tolQ* mutants had a PAGE LPS profile that was indistinguishable from the WT. The *wzm*, *rfbA*, *wbiF*, *wbiG* and *wbiI* mutants produced a similar LPS that lacked the O-antigen but had a lipid A/core oligosaccharide moiety that was indistinguishable from WT while the *waaC* and *waaF* mutants had an altered lipid A/core moiety, in agreement with the predicted heptosyl-transferase activity of the encoded enzymes that perform the first steps of the core oligosaccharide synthesis. Mass spectrometry analysis of the lipid A moieties suggested that none of the mutants had an altered lipid A structure and notably, that all mutants produced lipid A carrying the 4-amino-4-deoxy-L arabinose (Ara4N) modification that is known to confer PMB resistance in related species (34, 35, 39)(Fig. 4B; SI Appendix, Fig. S6).

**Figure 4.**
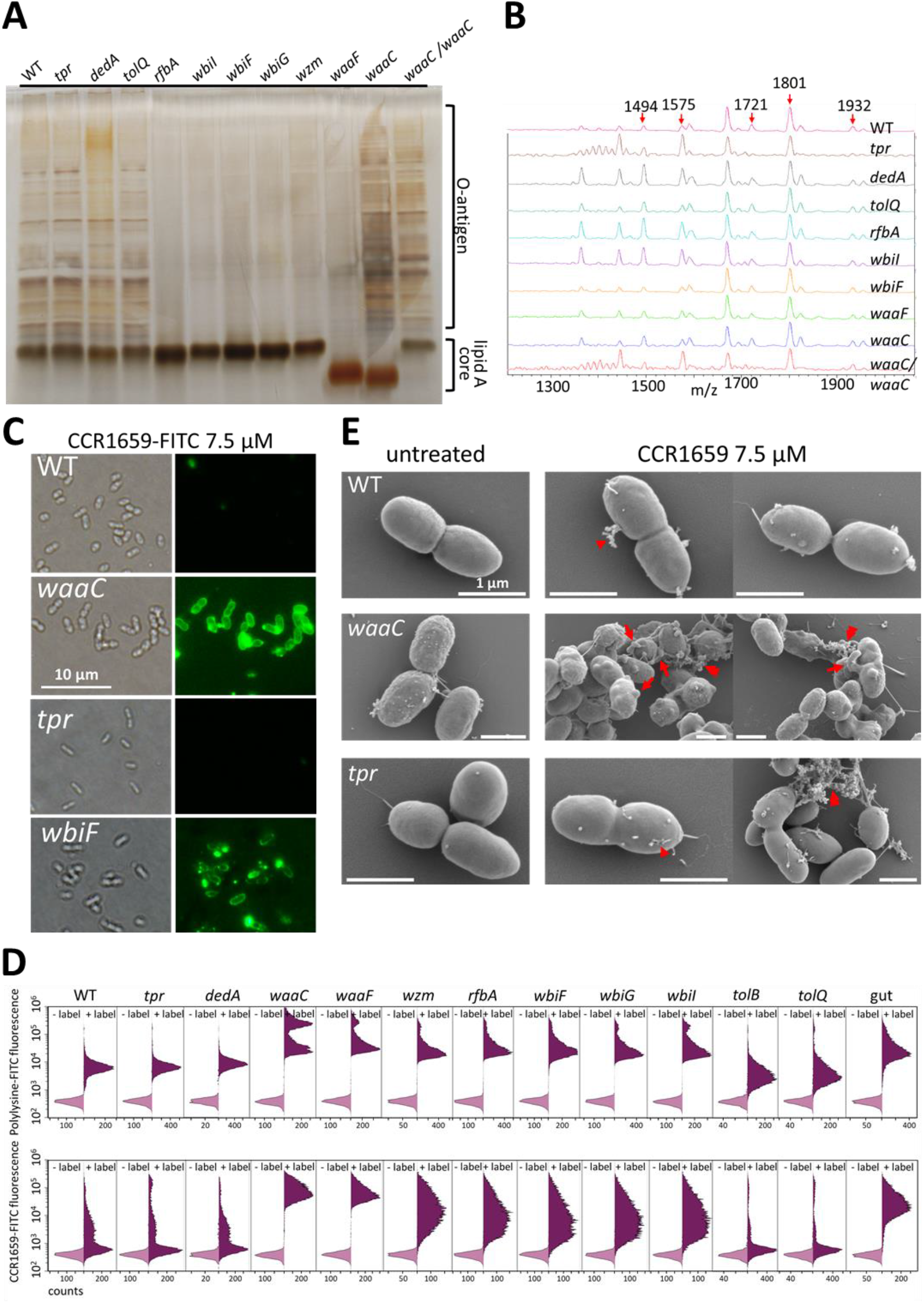
Surface properties of the AMP-sensitive mutants. (**A**) Polyacrylamide gel electrophoresis analysis of total LPS extracted from the indicated strains. The *waaC*/*waaC* strain is the complemented mutant. Despite the altered core in the *waaC* mutant, an O-antigen ladder is visible, that has a similar profile to the wild type, possibly corresponding to the O-antigen anchored on an intermediate lipid carrier. (**B**) MS analysis of the lipid A molecule present in the indicated mutants. Red arrows indicate the Ara4N carrying lipid A (Fig. S7). (**C**) Fluorescence microscopy of *C. insecticola* wild-type, *waaC*, *tpr* and *wbiF* cells treated with 50 µg/mL CCR1659-FITC. All images are at the same magnification and the scalebar is 10 µm. (**D**) Flow cytometry analysis of 50 µg/mL Polylysine-FITC (top) or 7.5 µM CCR1659-FITC binding by the indicated strains. Gut is bacteria directly isolated from the midgut crypts. Light purple histograms are control measurements without fluorescent label (-label); the dark purple histograms are in the presence of the fluorescent label. Note the presence of a double peak in the Polylysine-FITC treated mutants *waaC*, *waaF*, *wzm*, *rfbA*, *wbiFGI*, indicating of a heterogeneous bacterial population. (**E**) SEM micrographs of untreated *C. insecticola* wild type and *waaC* and *tpr* mutant untreated cells or treated with 7.5 µM CCR1659. Arrowheads indicate release of tiny amounts of cellular material in intact cells. Double arrowheads indicate cellular material released from lysed cells. Arrows indicate cell deformations. Scale bars are 1 µm for all images.

We assessed the steady-state outer membrane integrity of the mutants by NPN labeling and sensitivity to detergents (SI Appendix, Fig. S7). The *waaC* and *waaF* mutants had a higher NPN-derived fluorescence and slightly higher sensitivity to the non-ionic detergent Triton X100 and the cationic detergent CTAB than the WT, while the other mutants were similar to WT. The *tolB* and *tolQ* mutants on the other hand were more sensitive to the anionic detergent SDS than to other tested strains. Overall, this indicates that although the outer membrane in some mutants has a reduced robustness, the AMP sensitivity of the mutants is not a direct consequence of a generic membrane instability but of the deficiency of specific resistance mechanisms. The capacity of the bacterial envelope to bind membrane-disrupting AMPs is a parameter influencing AMP sensitivity. The *waaC* and *wbiF* LPS mutants showed indeed a strong labeling of their envelope with CCR1659-FITC, contrary to the WT that did not show any labelling (Fig. 4C). However, the *tpr* mutant was also not labeled. Therefore, we quantified the relative capacity of the envelope of all the mutants to bind membrane-disrupting AMPs by labeling the cells with the fluorescent polylysine-FITC peptide or CCR1659-FITC, followed by flow cytometry analysis (Fig. 4D). All the mutants with altered LPS (*waaC*, *waaF*, *wzm*, *rfbA*, *wbiF*, *wbiG* and *wbiI*) had a strongly enhanced labeling with both peptides indicating a more accessible cell surface for AMP binding. However, the *dedA* and *tpr* mutants displayed a peptide labeling that was identical to the WT while the *tolB* and *tolQ* mutants were even labelled less intensively. Thus, the LPS mutants might be more sensitive to the AMPs because of the higher accessibility of their membranes for interactions with AMPs but the sensitivity of the *dedA*, *tpr* and *tolBQ* mutants has to be explained by a different mechanism. Interestingly, crypt-colonizing *C. insecticola* bacteria have lost their O-antigen after establishing in the crypts (23) and thus have an LPS that is similar to the LPS of the *wzm*, *rfbA*, *wbiF*, *wbiG* and *wbiI* mutants. In agreement, bacteria isolated from the crypts are hypersensitive to PMB and the CCR1659 peptide (Fig. 3C, D) and they strongly bind polylysine-FITC and CCR1659-FITC (Fig. 4D).

To confirm that the set of mutants are affected in different pathways for AMP resistance, we created the *waaC*/*tpr*, *waaC*/*dedA* and *waaC*/*wbiF* double mutants. We reasoned that if genes are part of the same pathway, double mutants should not show an additive phenotype compared to the single mutants, while in case genes are in separate pathways, double mutants might display a more severe phenotype than single mutants. We found that the three double mutants were more sensitive than the corresponding single mutants to PMB and CCR1659 and bound more CCR1659-FITC (SI Appendix, Fig. S8), suggesting that indeed “*waaC* and *tpr*” or “*waaC* and *dedA*” or “*waaC* and *wbiF*” define different pathways to PMB resistance. The synthetic phenotype of the *waaC*/*wbiF* mutant further suggest that the LPS core and the O-antigen constitute two distinct barriers for AMPs to reach the membrane.

SEM of untreated WT and *tpr*, *dedA*, *tolB*, *waaC* and *wzm* mutants showed that the mutants affect the bacterial envelope in various ways (SI Appendix, Fig. S9). SEM of CCR1659-treated cells reveals that the response to the peptide in the *waaC* and *tpr* mutant is markedly different. In the *waaC* mutant, very strong membrane distortions are visible and frequent cell lysis, indicated by the cellular material released from cells. The *tpr* mutant on the other hand shows only minor modifications on the cell surface, similar to WT, although infrequent release of large amounts of cellular material was also observed (Fig. 4E). Collectively, the properties of the single and double mutants suggest that in *C. insecticola* different mechanisms contribute to AMP resistance.

### AMP resistance in *Caballeronia insecticola* is crucial for midgut colonization

Since the midgut crypts are the site of intensive AMP production, we next analyzed the capacity of the AMP sensitivity mutants to colonize the M4 midgut region of the *R. pedestris* midgut. As a preliminary test and to exclude that gut colonization phenotypes can be attributed to trivial reasons, we confirmed that each mutant has similar growth patterns as WT (SI Appendix, Fig. S10A) and is motile (SI Appendix, Fig. S10B) since motility is crucial for colonization of the M4 crypts (14). Analysis at 5 days post infection (dpi) of second instar nymphs showed that the 11 mutants had the capacity to colonize the crypts although they were to various extends less efficient than the WT. The WT had a 100% efficiency (n=10) and the number of bacteria per gut was consistently high (>10^7^ genome copies per gut). In contrast, the mutants displayed a large variability in colonization level between insect individuals, ranging from a wild-type colonization level for some individuals to a failure to establish in the crypts in other individuals (Fig. 5A). The *waaC* and *tpr* mutants were particularly affected in agreement with their strong AMP sensitivity. This intriguing probabilistic colonization of the gut by the mutants is reminiscent to stochastic colonization of the *Drosophila* gut by underperforming *Lactobacillus plantarum* strains while a strong colonizer strain had a 100% efficiency (40).

**Figure 5.**
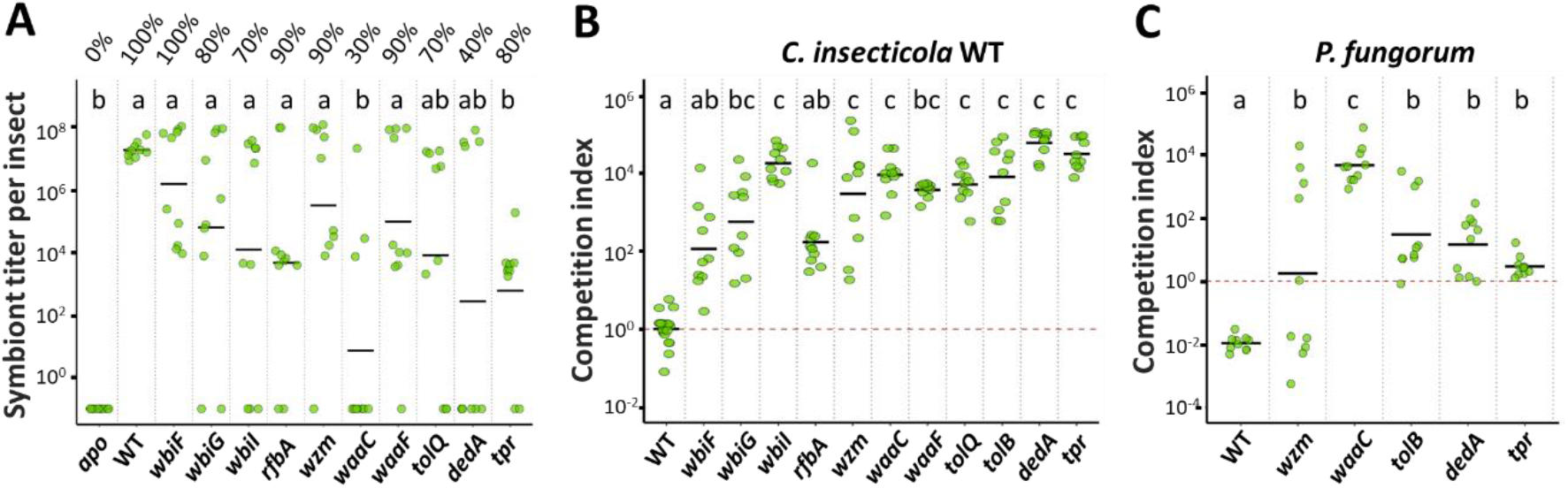
AMP-sensitive mutants are impaired in gut colonization. (**A**) Single-st ra in infections of *R. pedestris* second instar nymphs with *C. insecticola* WT or indicated mutants or no bacteria (*apo*). Colonization of the M4 crypt region was determined at 5 dpi by dissection and microscopy observation of the guts and symbiont titer determination by qPCR in M4 total DNA extracts. The % above the dot plots indicate the proportion of insects that showed colonization by microscopy observation (n=10). The qPCR measurements for each individ ual insect are indicated by green dots and the mean per mutant is indicated by a horizontal black line. (**B**) Co-infections of *R. pedestris* with an equal mix of RFP-labelled *C. insecticola* WT and indicated GFP-labelled WT or mutant strains. Relative abundance of the two strains in the M4 midgut regions at 5 dpi was determined by flow cytometry on dissected intestines. The competition index expresses for all samples the ratio of WT to the indicated mutant, corrected by the ratio of the inoculum, which was in all cases close to 1. Each dot represents the competition index in an individual and the mean per mutant is indicated by a horizontal black line (n=10). (**C**) Co-infections of *R. pedestris* with a 1:1 mix of GFP-labelled *P. fungorum* and indicated mScarlett-I-labelled WT or mutant *C. insecticola*. Relative abundance of the two strains in the M4 midgut regions at 5 dpi was determined by flow cytometry on dissected intestines. The competition index expresses for all samples the ratio of *P. fungorum* to the indicated mutant, corrected by the ratio of the inoculum, which was in all cases close to 1. Each dot represents the competition index in an individual and the mean per mutant is indicated by a horizontal black line (n=10). In all panels, different letters indicate statistically significa nt differences (*P*<0.05). Statistical significance was analyzed by Kruskal–Wallis test, Dunn *post hoc* test and Benjamini-Hochberg correction.

Next, we evaluated the fitness of the mutants in M4 colonization when they were in competition with WT. Insects were infected with fifty-fifty mixtures of RFP-marked WT and one of the mutants (or WT as a control) that were marked with GFP. The outcome of the competitions was analyzed at 5 dpi by fluorescence microscopy of dissected M4 midguts and flow cytometry quantification of their bacterial content (Fig. 5B). In the control competition, RFP- and GFP-marked WT were kept in balance. However, in competitions with the mutants, the WT nearly completely outcompeted each of them, confirming their reduced coloniza t ion capacity.

Finally, we also tested if the mutants have maintained or lost the capacity to outcompete a less efficient crypt colonizing species. We previously showed that *P. fungorum* can efficiently colonize the M4 crypts in the absence of competing strains but that it is outcompeted by *C. insecticola* when both strains are co-infecting the *R. pedestris* midgut (18). Here, the outcompetition of *P. fungorum* by *C. insecticola* WT in co-infection experiments was confirmed while *wzm*, *waaC*, *tolB*, *tpr* and *dedA* mutants were significantly less efficient in outcompeting *P. fungorum* (Fig. 5C). Thus, high AMP resistance in *C. insecticola* is an important factor contributing to the efficiency of this strain in occupying the *R. pedestris* gut.

## Discussion

The microbiota biogeography in the *R. pedestris* midgut shows a sharp divide between the anterior midgut, which has a highly variable, diverse and relatively low abundant microbiota, and the posterior midgut region that carries in striking contrast a dense mono-specific bacterial population that is strictly a *Caballeronia* species. The two principal findings from this work are that this posterior midgut region and the immediately adjacent anterior region is a highly challenging environment for bacteria because of the abundant presence of symbiosis-specific, membrane-damaging antimicrobial CCRs and that resistance to these AMPs is crucial for bacteria to colonize the crypts in the posterior midgut. Thus, we propose that the CCRs are new players, acting together with previously identified mechanisms (14–18), in the creation of the biogeography by eliminating sensitive bacteria.

The expression of the majority of the *CCR* genes is correlated with crypt coloniza t ion because they are expressed in the M4 crypt region of the midgut and they are frequently induced by bacterial colonization of the crypts. A few of them are also expressed in the upstream M3 midgut region, where they still may have a function related to crypt colonization, for example by preselecting bacterial species. Overall, their gene expression pattern, combined with their secretory nature suggesting that they are released in the lumen of the midgut, is consistent with a function of the CCRs in interacting with the bacterial community during midgut infection as well as during colonization.

Our analyses demonstrated that CCR peptides act through membrane interaction and damage, similarly to most AMPs produced by eukaryotic organisms (29, 41). AMPs damage membranes by first interacting with negative charges exposed on the membrane. In many Gram-negative bacteria, the negative charges carried by phosphate groups on the lipid A moiety of LPS are particularly important for this electrostatic interaction (27, 28). However, in *C. insecticola*, including in the AMP-sensitive mutants, these lipid A charges are converted into positive charges through the Ara4N modifications and therefore, the lipid A is likely not the target of AMPs in *C. insecticola*. We propose that the O-antigen and core oligosaccharide of LPS form a safeguard around the cell that limits the access of AMPs to their targets in the membrane. This hypothesis is consistent with the enhanced sensitivity and peptide-binding of mutants without O-antigen or LPS core. The direct targets of the AMPs are presently unknown but could be revealed by the analysis of the other genetic determinants of AMP resistance in *C. insecticola* identified here, including *tpr*, *dedA*, *tol-pal*, *tamAB*, *rpoE*, and hopanoid and phospholipid biosynthesis genes.

We conclude from our infection experiments that the reduced resilience to CCRs of *C. insecticola* mutants in different resistance mechanisms makes them less apt to colonize the midgut crypts. This correlates with the inability of strongly sensitive bacterial species to colonize the midgut (18). Presumably, AMP-resistance is critical during the initial infection stages, when a few hundred cells enter into the crypt region and this founder population subsequently multiplies rapidly, in two to three days, to a crypt-space-filling population of about 10^7^-10^8^ bacteria (14, 15). The surprisingly large diversity of CCR peptides, several of them already expressed in the M3 and M4 before the microbiota establishment, could be an adaptation to create a selective environment that restricts the type of bacteria from the anterior midgut microbiota that have a chance to establish in the M4 crypts and that favors optimal beneficial *Caballeronia* strains. Such a molecular filter of bacteria could arise from additive, synergistic or specific antimicrobial activities of CCR peptides towards distinct bacteria. Indeed, the tested CCRs have variable antimicrobial efficiency against different bacterial species and *C. insecticola* mutants. Recent insights from *Drosophila* and other models have changed the previous view on AMPs as generic, non-specific antimicrobials by the demonstration that they can display a degree of specificity and synergism. Accordingly, AMP repertoires in organisms dynamically evolve according to the diversity of microbes encountered in the natural environment (29, 41, 42). The hundreds of diverse CCR peptides might be an extreme example of such an evolutionary process.

On the other hand, once established in the crypts, the bacteria lose their O-antigen by an unknown mechanism (23), which renders them sensitive to the CCR peptides. This suggests a second function of the CCR peptide arsenal that could be related to the protection of the crypt epithelia and prevention of the bacteria breaching these epithelia. Indeed, in *R. pedestris* the crypt epithelium lack mucus or peritrophic protective layers and is therefore in direct contact with the microbiota (16). Additionally, the membrane fragilization of the crypt-bacteria by the CCR peptides could facilitate the retrieval of nutrients from the bacteria (43), suggesting that the insect tames the gut symbionts with the CCRs.

## Materials and Methods

### Bacterial strains and growth conditions

*Caballeronia insecticola* and *Paraburkholderia fungorum* strains were routinely cultured in Yeast-Glucose medium (YG: 5 g/L yeast extract; 1 g/L NaCl; 4 g/L glucose) at 28°C. *C. insecticola* RPE75 is a spontaneous rifampicin-resistant derivative of the wild-type strain RPE64 (18). *Sinorhizobium meliloti* Sm1021 and *Bacillus subtilis* Bs168 were grown in YEB medium (0.5 % beef extract, 0.1 % yeast extract, 0.5 % peptone, 0.5 % sucrose, 0.04 % MgSO_4_.7H_2_O, pH 7.5) at 28°C or LB medium (5 g/L yeast extract, 10 g/L tryptone, 5 g/L NaCl) at 37°C, respectively.

For standard molecular microbiology purposes, *Escherichia coli* strains DH5α, WM3064, MFDpir, S17-1λpir and their derivatives were grown in LB medium at 37°C. Growth of the Δ*dapA* MFDpir and WM3064 strains, which are auxotroph for diaminopimelic acid (DAP) synthesis, required the supplement of 300 µg/mL DAP to the medium. When appropriate, antibiotics were added to the medium in the following concentrations: 50 µg/mL kanamycin (Km) for *E. coli* and 30 µg/mL for *C. insecticola*; 25 µg/mL chloramphenicol (Cm); 30 µg/mL rifampicin (Rif); 100 µg/mL ampicillin (Amp); 500 µg/mL streptomycin (Sm) for *S. meliloti*. For solid agar plates, the media were supplemented with 1.5 % agar. For motility assays of *C. insecticola* derivatives, 5 µL of exponential phase cultures at OD_600nm_≈0.5 were injected in the centre of YG soft agar plates (0.3 % agar). Swimming motility of the tested strains was observed by the spreading of the bacterial growth from the inoculation point and was quantified by measuring the size of the formed halo using pictures of the plates, taken at regular time points after inoculation.

For growth inhibition assays, *C. insecticola* RPE75, *P. fungorum* JCM21562, *S. meliloti* Sm1021, *B. subtilis* Bs168 were grown in the minimal medium MM9 (40 mM MOPS; 20 mM KOH; 19.2 mM NH_4_Cl; 8.76 mM NaCl; 2 mM KH_2_PO_4_; 1 mM MgSO_4_.7H_2_O; 0.25 mM CaCl_2_.2H_2_O; 1 µg/mL biotin; 42 nM CoCl_2_; 38 µM FeCl_3_; 10 mM glucose) while *C. insecticola* and derived mutants were grown in the minimal medium MM (1 g/L KH_2_PO_4_; 2 g/L K_2_HPO_4_; 1 g/L (NH_4_)2SO_4_; 0.2 g/L NaCl; 0.1 g/L MgSO_4_.7H_2_O; 2.46 mg/L FeSO_4_.7H_2_O; 3.31 mg/L Na_2_EDTA.2H_2_O; 50 mg/L CaCl_2_.2H_2_O; 2 g/L glucose).

### Riptortus pedestris rearing

The bean bug *Riptortus pedestris* was originally collected from a soybean field in Tsukuba, Japan in 2007. For long-term maintenance of the insect line in the laboratory, they are reared in plastic boxes at 25°C under a long-day regimen (16 h light, 8 h dark) and fed with soybean seeds and distilled water containing 0.05 % ascorbic acid (DWA). Eggs are deposited on natural-fibre ropes present in the cages. Ropes carrying eggs are regularly collected and placed in a separate container. New-born hatchlings in these containers are collected daily and reared in fresh containers. At the second instar, a suspension of wild-type *C. insecticola* RPE64 cells at 10^7^ cfu/mL is added to the drinking water to keep the animals in the symbiotic state, which is optimal for reproduction (13).

### Riptortus pedestris colonization assays

Freshly hatched first instars were transferred into sterile Petri dishes. Two days after hatching, at the second larval stage, water was removed to make insects thirsty and facilitate the subsequent ingestion of administered bacteria. After overnight water-starvation, a bacterial suspension of the tested bacteria, grown in exponential phase at OD_600nm_≈0.5-0.7 and adjusted to 10^7^ cfu/mL in sterile distilled water, was provided for infection of the second instar nymphs insects. For co-inoculation experiments, two bacterial strains were mixed together at a one to one ratio, each adjusted before to 10^7^ cfu/mL. At three and five days post inoculation (dpi), insects, at the stage of the end of the second instar nymphs or the third instar, respectively, were dissected. Dissections were performed under binocular microscope in sterile PBS (137 mM NaCl, 8.1 mM Na_2_HPO_4_, 2.7 mM KCl, and 1.5 mM KH_2_PO_4_, pH 7.5) containing 0.01 % of Tween 20. The M4 region of the midgut was collected using fine forceps and assembled on glass slides for microscopy observations (Nikon Eclipse 80i). The colonization rate of the inoculated insects was estimated by detection of the presence or absence of fluorescent signal derived from colonizing bacteria expressing GFP or mScarlett-I fluorescent proteins. Merged fluorescence pictures were obtained with GIMP version 2.10.32. For quantification, M4 region samples were homogenized in PBS solution and bacteria in suspension were counted by plating on selective YG medium, by qPCR or by flow cytometry using the fluorescent tags to determine the relative abundance of the two inoculated strains. For each condition, 10 insects were tested within an experiment and each experiment was performed at least twice.

To quantify the symbionts in the M4 after infection of the insects with single strains, DNA was extracted from the dissected M4 by the QIAamp DNA Mini Kit (Qiagen), and real-time qPCR was performed using primers opcP-F and opcP-R (SI Appendix, Table S1), which target the *C. insecticola opcP* gene, the Fast Start Essential DNA Green Master qPCR Kit (Roche), and a LightCycler 96 instrument (Roche). The number of gut symbionts was calculated based on a calibration curve for the *opcP* gene consisting of ten-fold dilutions of 10^7^ to 10 copies of the PCR DNA fragment. Data were analysed with the instrument software and transferred to Microsoft Excel. Statistical analysis, using a Kruskal-Wallis test, Dunn *post hoc* test and Benjamini-Hochberg correction, was performed in R.

For co-inoculation experiments, the relative abundance of the two tested strains in competition was quantified by flow cytometry using a CytoFlex S instrument operated by CytExpert 2.4.0.28 software (Beckman Coulter). Gating by the forward-scatter (FSC) and side scatter (SSC) dot plot permitted to collect signals specifically derived from bacteria. Doublets were discarded using the SSC_Area-SSC_Height dot plot. GFP fluorescence was excited by a 488-nm laser and collected through a 525/40 nm band pass filter; RFP fluorescence was excited by a 561-nm laser and collected through a 610/20 nm band pass filter. Data acquisition for a total of 50,000-100,000 bacteria was performed for each sample. Thresholds for considering positive events for GFP and RFP were determined using non-fluorescent control bacteria. Data was treated by CytExpert and Microsoft Excel software. Statistical analysis, using a Kruskal-Wallis test, Dunn *post hoc* test and Benjamini-Hochberg correction, was performed in R.

### Transcriptome analysis

For transcriptome analysis of the midgut, insects were reared and infected as above. For samples of symbiotic insects (Sym), first day second instars were infected with *C. insecticola* RPE75. For uninfected insect samples (Apo), the animals were reared continuously in the absence of bacteria. Insects were harvested in the second instar at 1 dpi, 2 dpi and 3 dpi and in the fifth instar at 12 dpi. Apo insects were harvested at the same time points. Insects were dissected and the M3, M4B and M4 midgut regions were harvested in RNAiso Plus (Takara Bio). This resulted in 24 samples (Apo and Sym insects × 4 timepoints × 3 midgut regions). For each condition, samples of 24 insects in second instar and 12 insects in the fifth instar were pooled. Total RNA was extracted from the pooled samples with the RNeasy mini kit (Qiagen). mRN A purification and fragmentation and the preparation of cDNA libraries were conducted using TruSeq RNA Library Prep Kit v2 kit (Illumina). The cDNA libraries were sequenced by Illumina HiSeq-2000. The obtained RNA-seq data were analyzed by the bcl2fastq software (ver. 2-2.18.12, Illumina, Inc., San Diego, CA, USA) and FastQC (ver. 0.11.5) (https://www.bioinformatics.babraham.ac.uk/projects/fastqc/) to keep only high-quality data for *de novo* assembly and differential expression analysis. Raw read errors were corrected using Rcorrector (44) (K-mer-based method), followed by removing read pairs where at least one read had an unfixable error. Trim Galore (https://www.bioinformatics.babraham.ac.uk/projects/trim_galore/) was used to remove adaptor sequences, short reads, and low-quality reads. After the preprocessing, between 4 and 18 million paired-end reads remained per sample with a total of 412 million paired-end reads. The RNA-seq sequencing data were deposited in the Sequence Read Archive (SRA), BioProject accession no. PRJNA1006624.

The preprocessed reads were provided to Trinity software (45) for *de novo* assembly. Three methods were used to assess the quality of the Trinity assembly. First, assembly statistics were calculated using TrinityStats.pl script from Trinity software: the assembly contained 135,346 contigs (transcripts), N50 was 2541 bases and the total number of assembled bases in the contigs was 156,984,167. Second, the alignment rate of the preprocessed reads to the Trinity assembly was evaluated using Bowtie 2 (46), which resulted in an overall alignment rate of 98,61%. Third, the completeness of the *de novo* assembled transcriptome was assessed using BUSCO (47) with the OrthoDB v10 ‘Insecta’ and ‘Hemiptera’ reference databases which resulted in>97% and >94% of the Insecta and Hemiptera BUSCO genes identified as complete, respectively. In addition, transcripts encoding all the 97 reference CCR (Crypt-specific Cysteine Rich) peptides were identified in the *de novo* transcriptome using blast (48). The *de novo* assembled transcriptome was deposited in Transcriptome Shotgun Assembly (TSA), BioProject accession no. PRJNA1006624.

Coding regions in the Trinity *de novo* assembly were predicted using the TransDecoder tool (https://github.com/TransDecoder/TransDecoder). Then, the CCR peptides were predicted from this assembled and translated sequences using a Hidden Markov Model homology search against the 97 reference CCR peptides of *R. pedestris* (20) with HMMER software (49). The jackhammer application was used with the CCR reference sequences as input profile. In an iterative process by jackhammer, newly found CCR sequences are added to the input data set and the search was repeated. The number of iterations was 100 and additional iterations did not yield new sequences. Sequences with E-values of less than 10^-5^ were selected. The final output sequences were further filtered using custom scripts to remove those without a signal peptide and longer than the longest CCR peptide in the reference CCR database used for the prediction. The process yielded a total of 310 transcripts, including the reference data set, that were annotated as encoding CCR peptides.

The complete set of CCR peptides were aligned using Multalin (http://multalin.toulouse.inra.fr/multalin/) (50) and the alignment obtained was adjusted manually to keep only the well-aligned ones and to highlight the conserved cysteine residues. The adjusted alignment was used in WebLogo (51) to generate a graphical representation of the consensus sequence from the multiple sequence alignment.

Transcript and gene-expression levels of the transcripts encoding CCR peptides were estimated for each sample using RSEM software (52) based on alignment and transcript abundance estimation. The DEGs (Differentially Expressed Genes) were identified using edgeR (53) with FDR < 0.05 and log2(FC) ≥ 1 resulting in 305 (among 310) DEGs. The clustered heatmap, generated by MEV software ver. 4.8.0, of *CCR* genes against samples was based on the normalized FPKM (Fragments Per Kilobase of transcript per Million mapped reads) expression values matrix. FPKM values were transformed according to Value = [(Value) – Mean(Row)]/[Standard deviation(Row)] and clustered with hierarchical clustering using Pearson Correlation distance metric and average linkage (54). The expression strength of CCR genes relative to the overall transcriptome was estimated on the basis of the highest FPKM value among the 24 experimental conditions for each transcript and the ranking of transcripts according to these highest FPKM values.

The following antimicrobial peptide *in silico* predictor tools were used to predict the AMP activity potential of the 310 CCR peptide sequences: Collection of Anti-Microbial Peptides (CAMP_R4_) (55), using the Random Forest (RF), Support Vector Machine (SVM) and Artific ia l Neural Network (ANN) algorithms; AMPpredictor in dbAMP (56); iAMPpred (57); Antimicrobial Peptide Scanner v2 (AMPscanner) (58); Antimicrobial Peptide Prediction at AxPEP (59). A consensus AMP prediction was considered when 6 out of the 7 predictions were positive.

### Whole-mount *in situ* hybridization

Whole-mount *in situ* hybridization on the *R. pedestris* midgut was performed with digoxigenin (DIG)-labeled *CCR043* cRNA probe, using a method essentially as described (60). In brief, a DNA fragment of the *CCR0043* gene was amplified from M4 cDNA by PCR (SI Appendix, Table S1). The amplified cDNA fragment was cloned into the pT7Blue T-vector (Novagen). Gene-specific antisense or sense DIG-labeled cRNA probes were obtained by *in vitro* transcription with T7 polymerase using the Roche DIG RNA labeling kit (Roche Diagnostics) following the instructions of the supplier. Midguts of symbiotic and aposymbiotic insects were obtained as above and the tissues were fixed in 4 % paraformaldehyde, permeabilized with Proteinase K treatment and re-fixed. DIG-labeled cRNA probe hybridization was performed in hybridization buffer (50 % formamide, 5× SSC (0.75 M NaCl and 0.75 M Na-citrate, pH 7.4), 5× Denhardt’s solution (0.2 % bovine serum albumin, 0.2 % Ficoll, and 0.2 % polyvinylpyrrolidone), 25 mg/mL sonicated salmon sperm DNA, and 0.1 % Tween 20) at 61°C for 16 h. Detection of mRNA was performed with anti-DIG-alkaline-phosphatase-conjugated antibody (Roche Diagnostics) and the alkaline phosphatase substrate 5-bromo-4-chloro-3-indolyl phosphate (BCIP) and 4-nitroblue tetrazolium chloride (NBT) (Roche Diagnostics). After washing, the gut samples were mounted between slide and cover slip and observed by microscopy.

### Polymyxin B Tn-seq screening

An aliquot of an available *Himar1* **transposon mutant** Tn-seq library (32) was diluted to obtain a suspension of 2×10^8^ cfu/mL. For each tested condition, 100 µL of this dilution was inoculated into 20 mL of MM, supplemented with Rif and Km, to obtain an initial inoculum of 10^6^ cfu/mL (OD_600nm_≈0.0015). In the control growth condition, no Polymyxin B (PMB) was added. For the test conditions, the minimal media was supplemented with three different concentrations of PMB: 1.25 µM, 5 µM and 10 µM. These concentrations were chosen after a sensitivity assay and represent respectively half MIC, 1/4 MIC and 1/8 MIC of wild type *C. insecticola*. The resulting cultures were incubated at 28°C, with shaking at 180 rpm. When the cultures reached an OD_600nm_≈1, corresponding to approximately 9 to 10 generations of multiplication, bacteria were collected by centrifugation at 3300 rcf for 20 min at 4°C and the pellets were stored at - 20°C until DNA extraction. Each condition was performed in triplicates.

Genomic DNA extraction and processing to obtain Tn-seq sequencing libraries for Illumina sequencing was performed as described (32) using materials provided in SI Appendix, Table S1. The Tn-seq sequencing library samples were sequenced by an Illumina NextSeq 500 instrument with 2 x 75 paired-end run at the I2BC sequencing platform (CNRS Gif-sur-Yvette, France). The generated data were demultiplexed, trimmed, filtered and mapped as described (32). Tn-seq sequencing data were deposited in SRA, BioProject accession no. PRJNA890438.

Tn-seq sequencing data was handled by TRANSIT Version 3.2.0 (61) using software parameters as before and implementing statistics of the software (32). Each experimental condition with PMB treatment was compared to the control condition without PMB as a reference. The Integrative Genomics Viewer (IGV) software, an interactive tool for the visual exploration of genomic data, was used to visualize the number of insertions per insertion sites in specific genes of interest (62).

### Construction of *Caballeronia insecticola* RPE75 deletion mutants, fluorescent protein tagged strains and complementation strains

To create deletion mutants, regions of 400 bp flanking a gene of interest (Up and Down regions) were obtained by gene synthesis (GenScript) as a fused 800 bp fragment (SI Appendix, Table S1) and cloned into the *Sma*I/*Hind*III or *EcoR*I/*Hind*III sites of the pK18mob*sacB* vector. The constructs were introduced in *E. coli* WM3064. Plasmids were introduced into *C. insecticola* RPE75 by bi-parental mating, resulting in first recombinant clones with the plasmids integrated in the genome. PCR with gene specific primers (SI Appendix, Table S1) and subsequent sequencing confirmed insertions at the appropriate chromosomal positions. To obtain second recombinants in which the gene of interest has been deleted, bacterial cultures of first recombinants were plated on YG containing 10 % sucrose and Rif to counter-select the *sacB* gene located on the pK18mob*sacB* plasmid. Candidate deletion mutants were verified by colony PCR (SI Appendix, Table S1).

GFP- or mScarlett-I-tagged strains of *C. insecticola* wild type and mutants were created by introducing a Tn7-GFP or Tn7-Scarlet transposon using tri-parental mating with the Tn7-GFP donor strain WM3064.pURR25 or the Tn7-Scarlet donor strain S17-1λpir.pMRE-Tn7-135, the helper strain WM3064.pUX-BF13 and *C. insecticola* wild type or derivatives as acceptor strains.

For the construction of complemented strains, the plasmid pME6000 was digested by *Hind*III and the PCR-amplified gene of interest was inserted by Gibson cloning (NEB). For this, DNA fragments containing about 350-400 bp of upstream non-coding sequence, the open reading frame and 300-350 bp of downstream non-coding sequence of a corresponding gene were amplified with gene-specific primers (SI Appendix, Table S1). PCR products were used as template for second PCR amplifications with Gibson-compatible primers (SI Appendix, Table S1). The constructed plasmids, cloned in DH5α, were introduced in *E. coli* WM3064 and transferred to the corresponding mutant by bi-parental mating.

### Antimicrobial peptide activity assays

Precultures of tested strains were grown in MM9 or MM medium. Overnight grown cultures were diluted to an OD_600nm_=0.3 in fresh medium and grown until they reached OD_600nm_≈1. The cells were pelleted by centrifugation, resuspended in fresh medium and diluted to OD_600nm_=0.05. For testing crypt-colonizing *C. insecticola*, 100 insects at 5 dpi were dissected, the M4 region collected and the bacteria extracted as described above. The obtained suspensions were diluted to OD_600nm_=0.05 in MM. These cell suspensions were dispatched in a 96-well plate, one column per tested strain and at 150 µL per we **l**, except for the we **l**s of the first row, which contained 300 µL. Peptides used in this study are listed in SI Appendix, Table S1. The CCR peptides were selected on the basis of the following criteria: 1) consistent prediction of AMP activity; 2) diversity of expression patterns, including peptides expressed in apo and/or sym insects and in the M3, M4B and/or M4, with high or medium expression levels; 3) favouring smaller peptides to increase feasibility of successful peptide synthesis; 4) successful synthesis by commercial suppliers (synthesis attempts for several peptides failed). AMPs, dissolved in water, were added to the first row to reach a maximal tested final concentration. Two-fold serial dilutions in the subsequent rows were obtained by the serial transfer of 150 µL to the next row and mixing by pipetting up and down. No peptide was added to the last row of the 96-well plate. This setup resulted in the testing of seven peptide concentrations forming a two-fold concentration series in the range of 120 µM to 0.5 µM and a control sample without peptide. The 96-well plates were incubated in a SPECTROstar Nano plate incubator (BMG Labtech). The growth of the cultures in the wells was monitored by measuring the OD_600nm_ and data points were collected every hour for 48 h. Plates were incubated at 28°C with double orbital shaking at 200 rpm. Data and growth curves were analyzed using Microsoft Excel. For the comparison of *C. insecticola*, *P. fungorum*, *S. meliloti* and *B. subtilis*, the minimal peptide concentration of growth inhibition was determined. For the comparison of *C. insecticola* wild type and derived mutants, the minimal concentration was determined at which growth was diminished compared to the untreated control. The assays were performed in biological triplicates for all peptides.

To determine bacterial survival after peptide treatment by cfu counting, *S. meliloti* was grown in MM9 until OD_600nm_<1. Bacteria were resuspended in fresh medium to OD_600nm_=0.01. CCR1659 peptide, Proteinase K-inactivated CCR1659 or PMB were added to the suspension to a final concentration of 25 µM. To inactivate peptide CCR1659, a peptide stock at 500 µM was treated with 0.1 mg/mL Proteinase K for 2h at 37°C. The treatments were made in the absence or presence of 5 mM CaCl_2_. Suspensions were incubated at 28°C and 10 µL samples were withdrawn at various time-intervals. 5 µL of ten-fold dilution series of these samples were spotted on plates with YEB medium for cfu counting. The assays were performed in triplicates.

### Membrane stability measurements

For evaluation of the membrane permeabilization of *S. meliloti* in response to CCR1659 or PMB, bacteria were grown in MM9 medium until OD_600nm_<1. Bacteria were resuspended to OD_600nm_=0.1 in fresh MM9, containing either 10 µM 1-N-phenylnaphthylamine (NPN) (63, 64) or 1 µg/mL Propidium Iodide (PI). Suspensions were dispatched per 100 µL in a black 96-well plate with transparent bottom (Falcon). CCR1659 peptide or PMB were added to the suspensions to a final concentration of 10 µM; control treatments were without peptide. All treatments were made in the absence or presence of 5 mM CaCl_2_. Fluorescence was measured at 28°C every 2 min for a total of 100 min incubation in an infinite M1000 PRO fluorescence plate reader (Tecan), operated with Tecan i-control, version 1.11.1.0 software. NPN fluorescence was excited at 340 nm and emission detection was at 420 nm. PI fluorescence was excited at 535 nm and emission detection was at 617 nm. Data was exported to Microsoft Excel for analysis.

Membrane stability properties in *C. insecticola* wild type and mutants were assessed by NPN fluorescence and sensitivity to detergents SDS, Triton X-100 and CTAB. The strains were grown in MM medium until OD_600nm_<1. For NPN fluorescence measurements, bacteria were resuspended to OD_600nm_=0.5 in fresh MM, containing 10 µM NPN. Fluorescence was measured as above after 10 min incubation in the NPN-containing medium. Detergent sensitivity assays were performed using maximum detergent concentrations of 0.02 % for SDS, 0.05 % for Triton X-100 or 0.001 % for CTAB in the microplate two-fold-dilution setup as above for the AMP sensitivity assays.

### Peptide binding to bacterial cells

Precultures of *S. meliloti* or *C. insecticola* strains were grown in MM9 or MM medium, respectively. Overnight grown cultures were diluted to an OD_600nm_=0.3 in fresh medium and grown until they reached OD600nm≈1. The ce **l**s were pe **l**eted by centrifugation, resuspended in fresh medium and diluted to OD_600nm_=0.1. Crypt-colonizing bacteria were obtained by dissecting 100 insects at 5 dpi, collecting the M4 region and extracting the bacteria by homogenizing the M4 tissues as described above. The obtained suspensions were diluted to OD_600nm_=0.1 in MM. FITC-CCR1659 or Poly-L-lysine-FITC (SI Appendix, Table S1) were added to a final concentration of respectively 1.5 µM or 10 µg/mL for *S. meliloti* and of 7.5 µM or 50 µg/mL for *C. insecticola* strains. Labelling of cells was analysed immediately after addition of the peptides by microscopy and flow cytometry. Labelled bacteria were spotted on glass slides covered with an agar pad for fluorescence microscopy observations (Nikon Eclipse 80i). Labelling was quantified by flow cytometry using a CytoFlex S instrument operated by CytExpert 2.4.0.28 software (Beckman Coulter). Gating by the forward-scatter (FSC) and side scatter (SSC) dot plot permitted to collect signals specifically derived from bacteria. Doublets were discarded using the SSC_Area-SSC_Height dot plot. FITC fluorescence was excited by a 488 nm laser and collected through a 525/40 nm band pass filter. Data acquisition for a total of 20.000 bacteria was performed for each sample. Control samples without addition of FITC-labelled peptide were used to determine the background signal of the bacteria. Data was treated by CytExpert and Microsoft Excel software.

### Scanning electron microscopy

*S. meliloti* was grown in MM9 and *C. insecticola* wild type and mutants in MM. Bacteria in exponential-phase (OD_600nm_<1) were resuspended in fresh medium to OD_600nm_=0.2. One mL of *S. meliloti* suspensions were treated with 1.5 µM CCR1659 or 3.6 µM PMB or without peptide. One mL of *C. insecticola* suspensions were treated with 1.5 M CCR1659 or without peptide. After 1 h of incubation at room temperature, bacteria were sedimented by soft centrifuga t ion (2 min, 2000 g), the bulk of the supernatant was removed and the remaining 50 µL was directly fixed in 2 mL 2 % glutaraldehyde buffered with sodium cacodylate 0.2 M, 2 h at room temperature then overnight at 4°C on glass slides. Samples attached to the glass slides were rinsed 10 min in 0.2 M sodium cacodylate buffer, dehydrated in successive baths of ethanol (50, 70, 90, 100, and anhydrous 100 %), and then dried using a Leica EM300 critical point apparatus with slow 20 exchange cycles, with a 2 min delay. Samples were mounted on aluminum stubs with adhesive carbon and coated with 6 nm of Au/Pd using a Quorum SC7620, 50 Pa of Ar, 180 s of sputtering at 3.5 mA. Samples were observed using the SE detector of a FEG SEM Hitachi SU5000, 2 KeV, 30 spot size, 5 mm working distance (facilities located on the MIMA2 platform, INRAE, Jouy-en-Josas, France; https://doi.org/10.15454/1.5572348210007727E12). Scanning electron microscopy (SEM) images were analyzed with FIJI software and calibrated using the printed scale bar on the image during the acquisitions.

### LPS and lipid A extraction and analysis

Bacteria were grown in 100 mL YG at 28°C until OD_600nm_≈2 and LPSs were isolated by the phenol/water method of Westphal and Jann (65). Briefly, the wet pellet of bacteria was stirred in 30 mL 50% aqueous phenol at 65°C for 15 min, insoluble material was removed from the cooled water phase by centrifugation and the clear extract was dialyzed under running tap water until free of phenol, then dialyzed against distilled water. The samples were subsequently lyophilized.

LPS sample analysis by SDS-PAGE was performed on 15 % polyacrylamide separation gels layered with 4 % polyacrylamide stacking gel, using standard electrophoresis buffers and migration conditions (66). About 0.5 µg LPS per sample was loaded. Gels were stained with the silver nitrate method essentially as described before (67).

Lipid A was prepared by the triethylamine-citrate method (68). Briefly, the LPS samples were suspended at a concentration of 10 µg/µl in a 0.01 M triethylamine-citrate solution (1:1 molar ratio, pH 3.6) and heated for 1 h at 100°C. The samples were then lyophilized and suspended in methanol. After centrifugation (7000× g for 10 min at 4°C), lipid A fractions were extracted with a mixture of chloroform: methanol: water (3:1.5:0.25, v:v:v) at a concentration of 10 µg/µL.

The molecular species present in this preparation were analyzed using an AXIMA performance (Shimadzu Biotech) matrix-assisted laser desorption ionization–time of flight (MALDI-TOF) mass spectrometer. A suspension of lipid A (1 µg/µL) in chloroform: methanol: water (3:1.5:0.25, v:v:v), 1 µL was deposited on the target mixed with 1 µL of a gentisic acid (2,5-dihydroxybenzoic acid) matrix (DHB from Fluka) suspended at 10 µg/µL in the same solvent, and dried. Analyte ions were desorbed from the matrix with pulses from a 337 nm nitrogen laser. Spectra were obtained in the negative-ion mode at 20 kV, with the linear detector. Mass calibration was performed with a *peptide mass standards* kit (*AB SCIEX) or with a* purified and structurally characterized LPS sample from *Bordetella pertussis*.

## Data availability

RNA-seq sequencing data are available in the Sequence Read Archive (SRA), BioProject accession no. PRJNA1006624. The *de novo* assembled transcriptome was deposited in the Transcriptome Shotgun Assembly (TSA), BioProject accession no. PRJNA1006624. Tn-seq sequencing data were deposited in SRA, BioProject accession no. PRJNA890438.

## Acknowledgments

We are grateful to Olga Soutourina (University Paris-Saclay, France) and Yu Matsuura (University of the Ryukyus, Japan) for critical reading of the manuscript and constructive comments. This work benefited from financial support by Saclay Plant Sciences-SPS, by the ANR grant ANR-19-CE20-0007, and by a JSPS-CNRS Bilateral Open Partnership Joint Research Project (18KK0211) and a CNRS International Research Project to Y.K. and P.M. Y.K. was supported by the MEXT KAKENHI (21K18241, 22H05068). J.L., G.L. and R.J. were supported by Ph.D. fellowships from the French Ministry of Higher Education, Research, and Innovation and A.B. benefited from a PhD contract in the frame of the CNRS 80|PRIME – 2021 program. This work was supported by JSPS Research Fellowships for Young Scientist to S.J. (21F21090), K.I. (22KJ0057) and T.O (14J03996, 20170267 and 19J01106). Tn-seq sequencing and data treatment were performed by the I2BC high-throughput sequencing facility, supported by France Génomique (funded by the French National Program Investissement d’Avenir ANR-10-INBS-09). This work has benefited from the facilities and expertise of MIMA2 (Université Paris-Saclay, INRAE, AgroParisTech, 78350, Jouy-en-Josas, France).

## Author contributions

P.M. and Y.K. designed the study, planned the experiments and supervised the project. T.O., R.F, A.B. and B.A. performed transcriptome analysis. J.L., R.J., D.N. performed Tn-seq. J.L., R.J. and T.T. made mutants. P.M. and E.B. performed in vitro peptide activity assays. S.J. and K.I. provided strains. V.C. performed SEM. L.A. and P.T. performed LPS characterizat io n. J.L., G.L., A.Y., R.C and T.O. performed insect experiments. J.L., G.L., R.J, A.B., T.O., Y.K, and P.M. analyzed data. P.M. wrote the manuscript with input from Y.K. All authors provided critical feedback and helped to shape the manuscript.

## Notes

### Competing Interest Statement

The authors have declared no competing interest.

### Summary of Updates

Abstract, the main text and its organization as well as figure 1 have been modified. There are no modifictions in the presented data.

