## Supplemental figures for "Hundreds of antimicrobial peptides create a selective barrier for insect gut symbionts"

**This PDF file includes:**

Figs. S1 to S10

**Other Supplementary Materials for this manuscript include the following:**

Table S1
Data S1 and S2

**Fig. S1.**

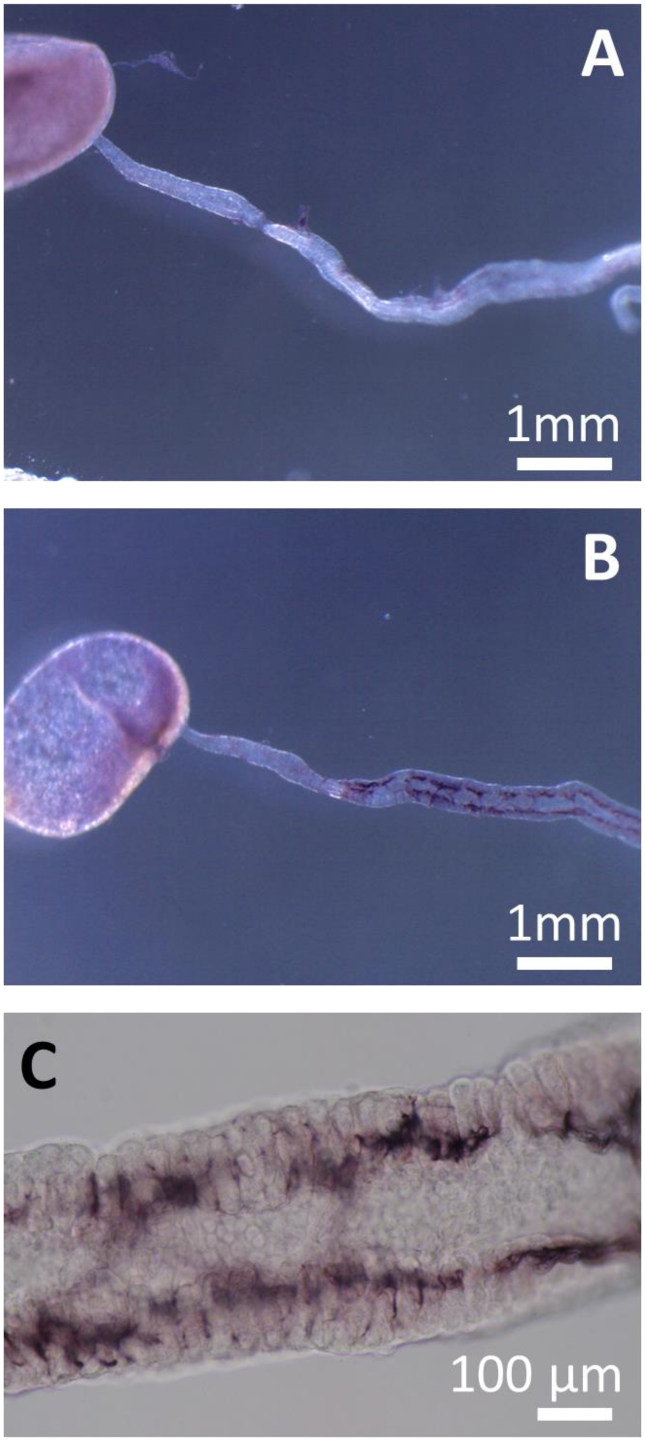


**Whole-mount *in situ* hybridization controls.** (**A**) Hybridization of a 3 dpi midgut with a CCR0043 sense probe revealing the absence of signal. (**B**) Hybridization of an aposymbiotic midgut with a CCR0043 antisense probe revealing a signal at the base of crypts. (**C**) Detail of crypts as in panel B.

**Fig. S2.**

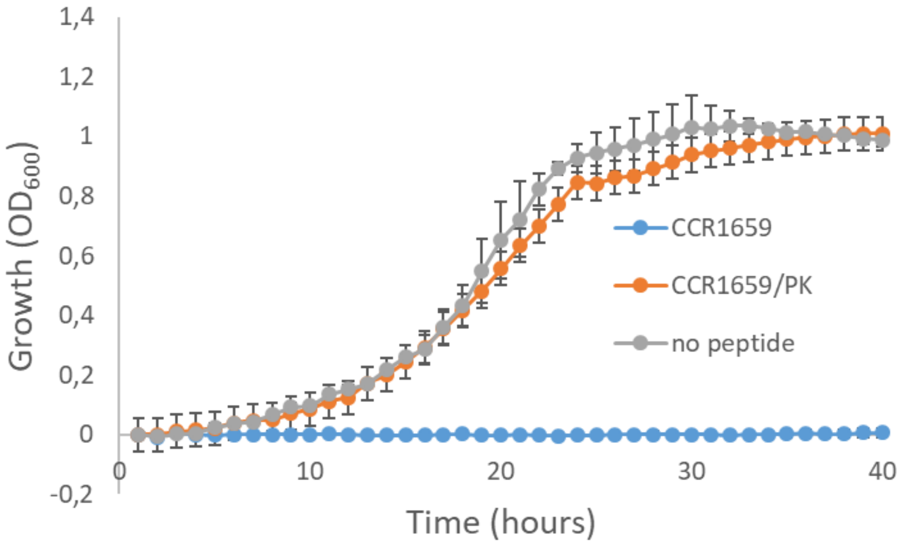


**Suppression of CCR1659 activity by Proteinase K treatment.** Growth curves of *S. meliloti* control or in the presence of 18 µM CCR1659 or an equivalent amount of CCR1659 treated prior with Proteinase K (PK). A peptide stock at 500 µM was treated with 0.1 mg/mL Proteinase K for 2h at 37°C before use in the growth assay. The control samples (no peptide) were grown in the presence of a similar amount of Proteinase K (2.5 µg/mL). Error bars are standard deviations (n=3).

**Fig. S3.**

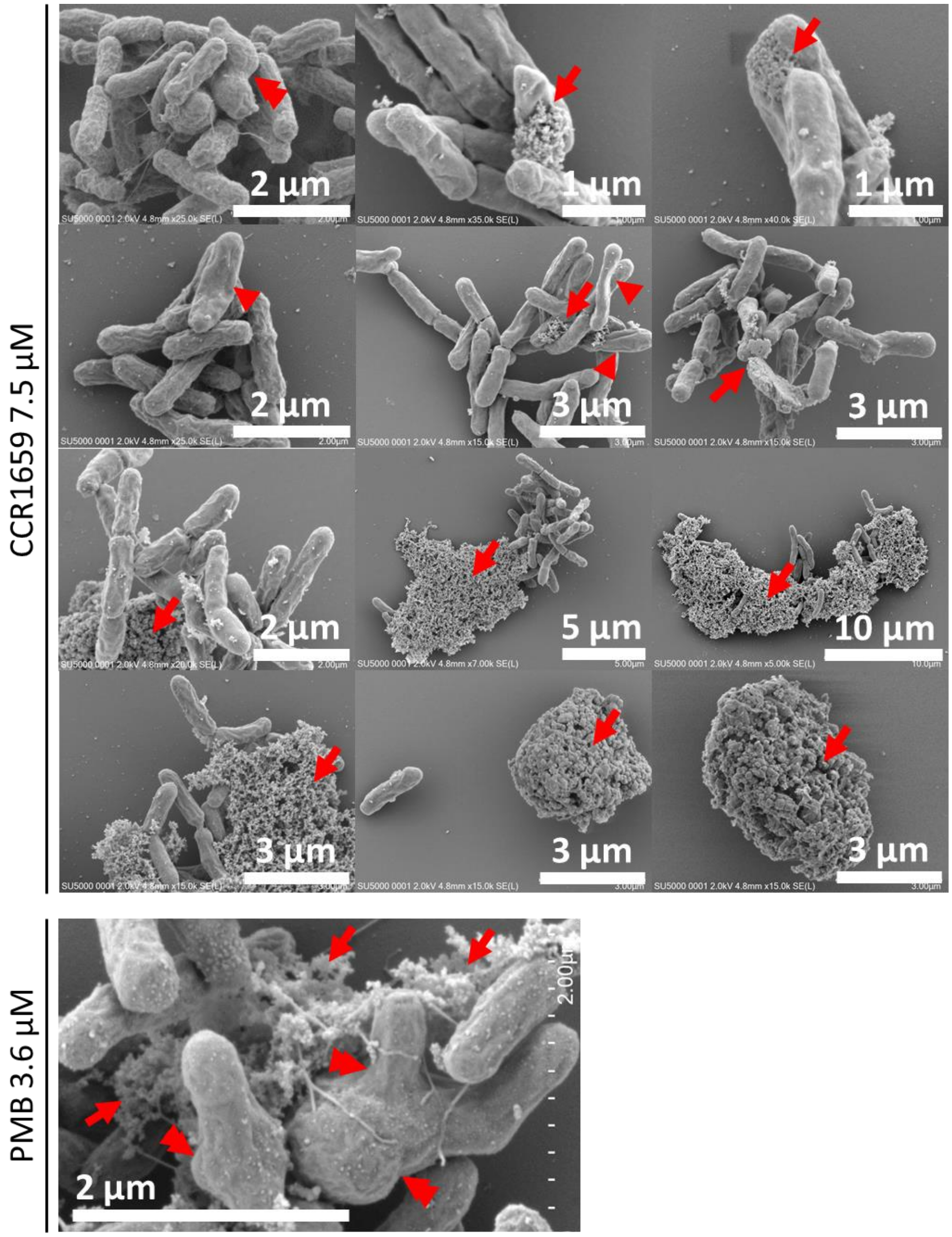


**SEM micrographs *S. meliloti* treated with CCR1659 or PMB.** Cells were treated with 7.5 µM CCR1659 or with 3.6 µM PMB. The arrows indicate cellular material released from cells. Arrowheads indicate cells with lost turgor. The double arrowheads indicate morphologically aberrant cells.

**Fig. S4.**

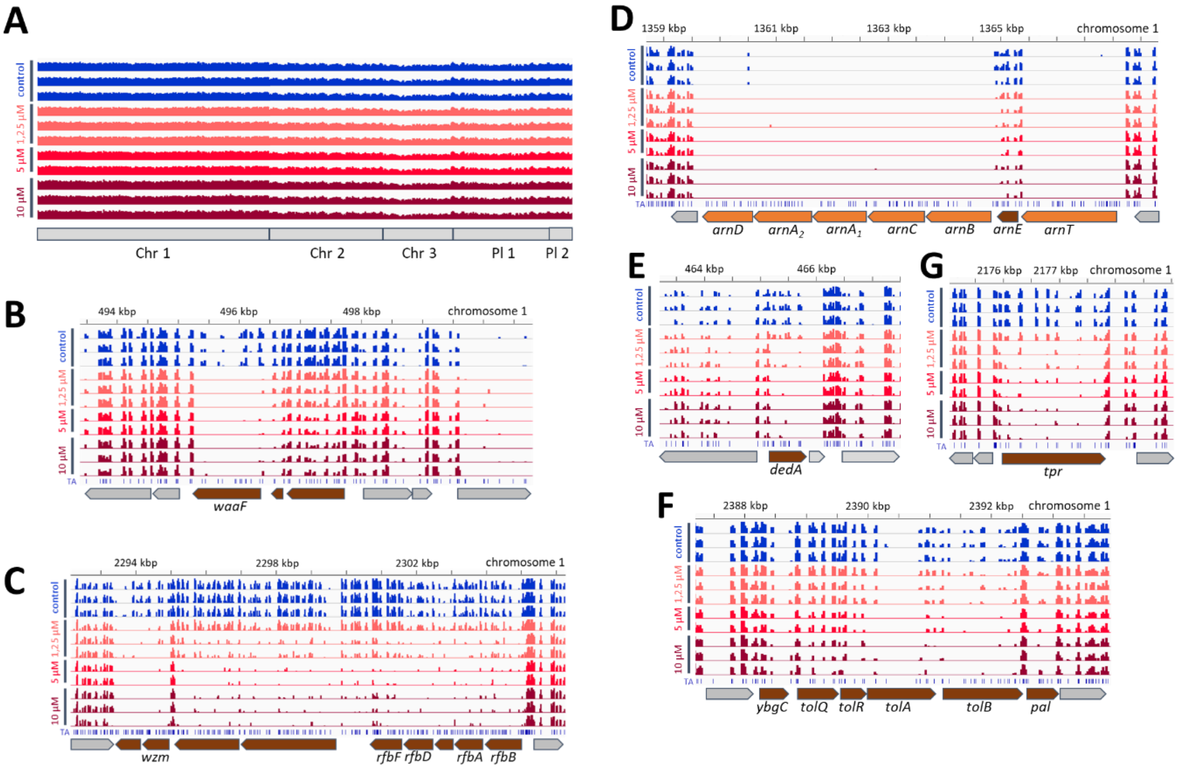


**IGV views of Tn-seq sequencing data of *Caballeronia insecticola* genome regions.** (**A**) Whole genome view showing the reproducible profiles in the control and PMB conditions. (**B-F**) Views of selected regions corresponding to the genes that were further characterized in this study.

**Fig. S5.**

| minimal concentration of growth inhibition (µM) | WT | *waaC* | *waaC/waaC* | *dedA/pME* | *dedA/dedA* | *tpr/pME* | *tpr/tpr* | *wbiF/pME* | *wbiF/wbiFG* | *wbiG/pME* | *wbiG/wbiFG* |
|---|---|---|---|---|---|---|---|---|---|---|---|
| PMB | 37,5 | 1,2 | 37,5 | 2,3 | 37,5 | 1,2 | 37,5 | 1,2 | 37,5 | 0,6 | 37,5 |
| COL | 43,3 | 1,35 | 43,3 | 5,4 | 43,3 | 2,7 | 43,3 | 2,7 | 43,3 | 2,7 | 43,3 |

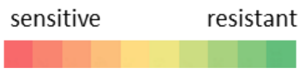


**Complementation of PMB sensitivity.** Minimal concentrations of growth inhibition of the indicated wild-type, mutant and complemented mutant strains by PMB and COL. Minimal concentrations are indicated in µM. Complementing genes were introduced on the plasmid pME6000. pME indicates the empty plasmid pME6000.

**Fig. S6.**

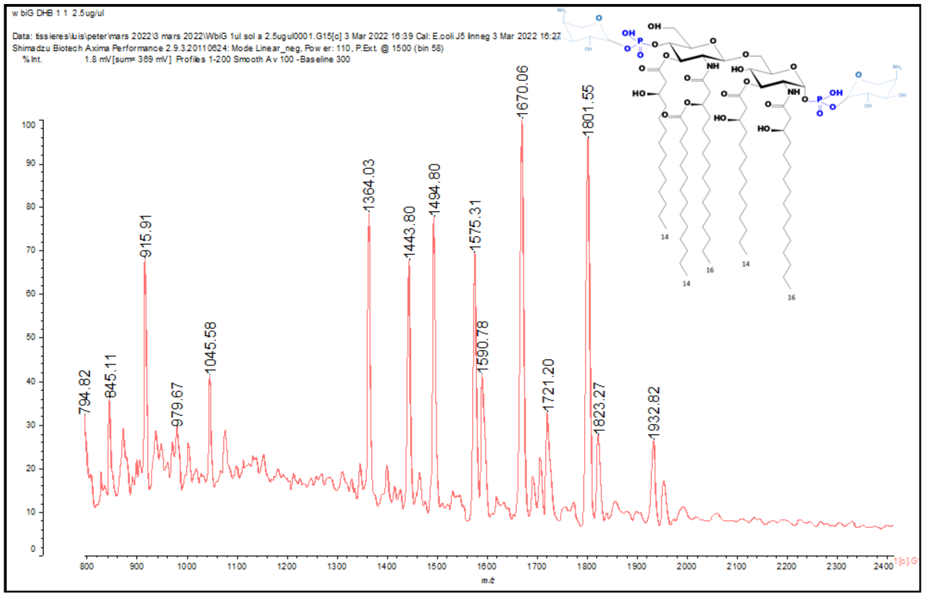


Peak 1364.03 = 2.GlcN + 1.P + 1.C14OH + 2.C16OH + 1.C14
Peak 1443.8 = 2.GlcN + 2.P + 1.C14OH + 2.C16OH + 1.C14
Peak 1494.8 = 2.GlcN + 1.P + 1.C14OH + 2.C16OH + 1.C14 + 1.AraN
Peak 1575.31 = 2.GlcN + 2.P + 1.C14OH + 2.C16OH + 1.C14 + 1.AraN
Peak 1590.78 = 2.GlcN + 1.P + 2.C14OH + 2.C16OH + 1.C14
Peak 1670.06 = 2.GlcN + 2.P + 2.C14OH + 2.C16OH + 1.C14
Peak 1721.0 = 2.GlcN + 1.P + 2.C14OH + 2.C16OH + 1.C14 + 1.AraN
Peak 1801.55 = 2.GlcN + 1.P + 2.C14OH + 2.C16OH + 2.C14
Peak 1801.55 = 2.GlcN + 2.P + 2.C14OH + 2.C16OH + 1.C14 + 1.AraN
Peak 1932.82= 2.GlcN + 2.P + 2.C14OH + 2.C16OH + 1.C14 + 2.AraN

**MS profile of *C. insecticola* lipid A.** The MS spectrum is shown for the lipid A molecule of *C. insecticola*. The masses of the peaks are indicated and their corresponding chemical composition is listed below. The inset shows the structure of the lipid A molecule with m/s= 1932,82 with its two Ara4N modifications shown in light blue.

**Fig. S7.**

| | WT | *tpr* | *dedA* | *tolB* | *tolQ* | *waaC* | *waaF* | *rfbA* | *wbiF* | *wbiG* | *wbiI* | *wzm* |
|---|---|---|---|---|---|---|---|---|---|---|---|---|
| NPN (au) | 1079 | 1231 | 1387 | 1142 | 1206 | 2407 | 2569 | 1479 | 1528 | 1462 | 863 | 1163 |
| SDS (%) | 0,02 | 0,02 | 0,02 | 0,01 | 0,01 | 0,02 | 0,02 | 0,02 | 0,02 | 0,02 | 0,02 | 0,02 |
| Trtion X100 (%) | 0,1 | 0,1 | 0,1 | 0,1 | 0,1 | 0,025 | 0,025 | 0,1 | 0,1 | 0,1 | 0,1 | 0,1 |
| CTAB (%) | 0,00025 | 0,00025 | 0,00025 | 0,00025 | 0,00025 | 0,000125 | 0,000125 | 0,00025 | 0,00025 | 0,00025 | 0,00025 | 0,00025 |

**Membrane integrity.** Steady state membrane integrity was measured in the wild type (RPE75) and indicated mutant strains, grown in MM medium in exponential growth phase, by NPN fluorescence (au, arbitrary units of fluorescence) and sensitivity to anionic (SDS), neutral (Triton X100) or cationic (CTAB) detergents. The indicated sensitivities (% detergent) indicate the minimal concentration to inhibit or diminish growth.

**Fig. S8.**

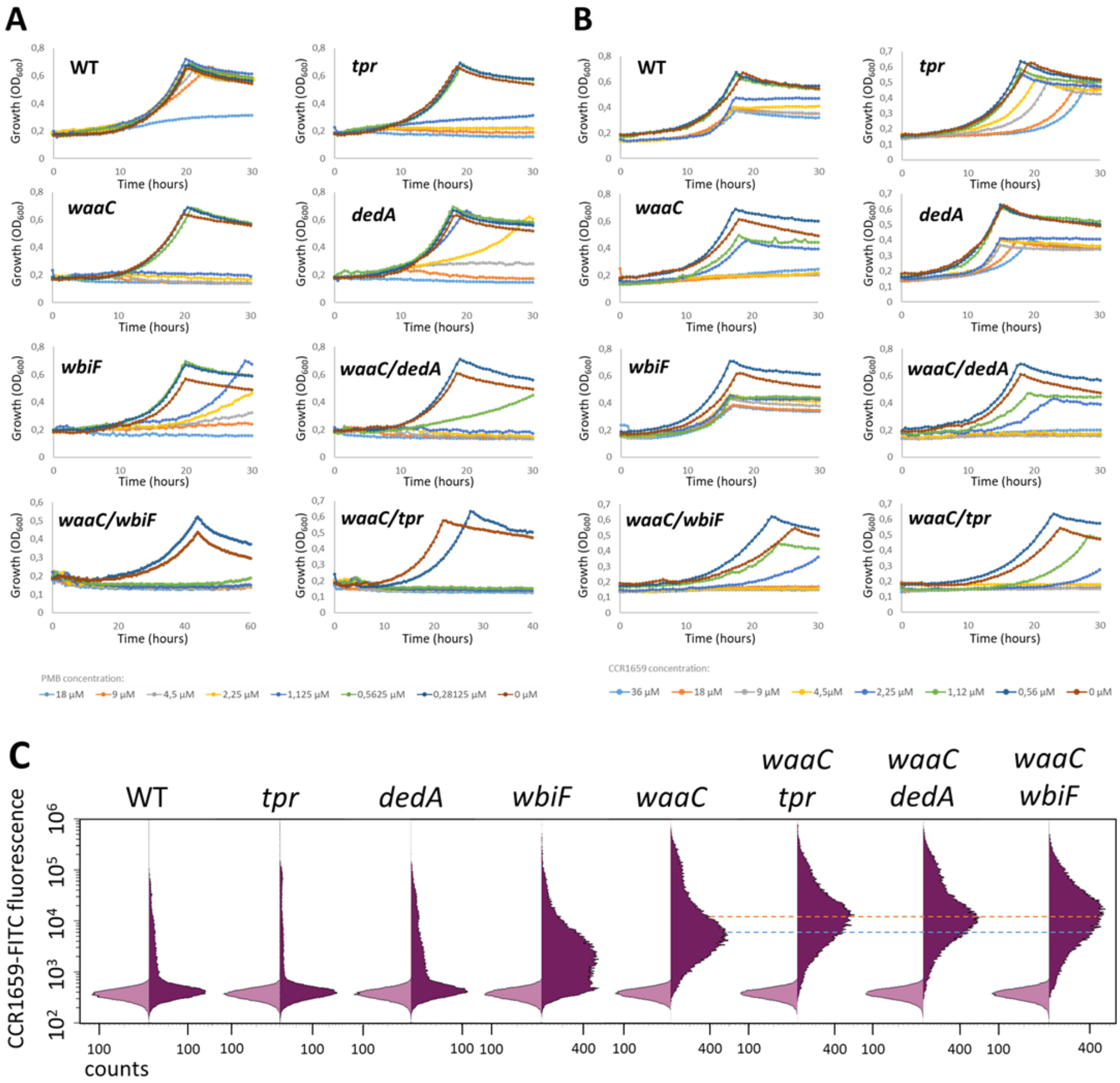


**Synthetic phenotype in double mutants.** (**A**) Growth inhibition of *C. insecticola* wild type, single mutants and double mutants by different concentrations of PMB. (**B**) Growth inhibition of *C. insecticola* wild type, single mutants and double mutants by different concentrations of CCR1659. (**C**) Flow cytometry analysis of 7.5 µM CCR1659-FITC binding by *C. insecticola* wild type, single mutants and double mutants. Light purple histograms are control measurements without fluorescent label; the dark purple histograms are in the presence of the fluorescent label.

**Fig. S9.**

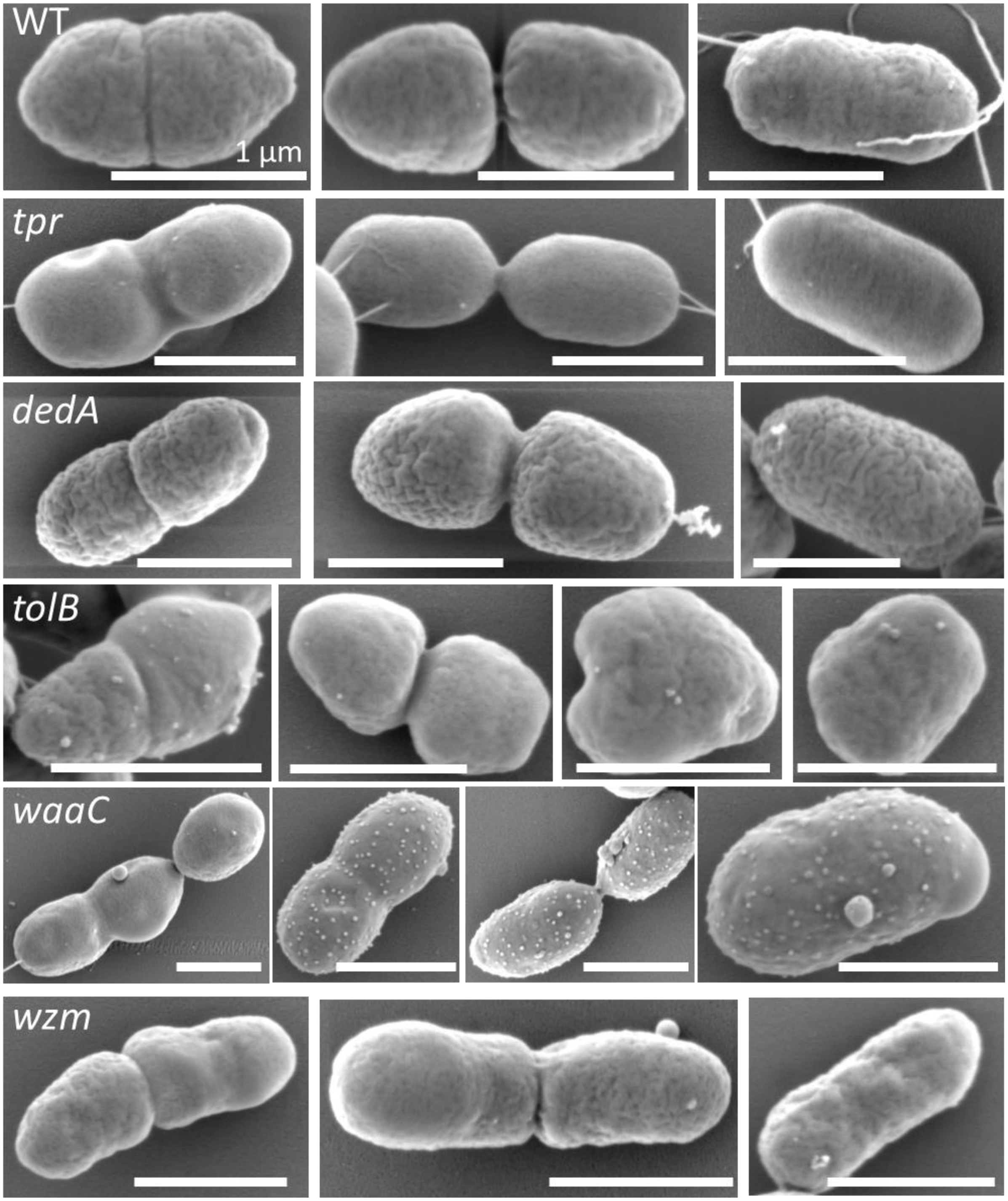


**AMP-sensitivity mutants have altered envelope and morphology properites.** SEM micrographs of untreated *C. caballeronia* wild-type and indicated mutant cells. Scale bars are 1 µm.

**Fig. S10.**

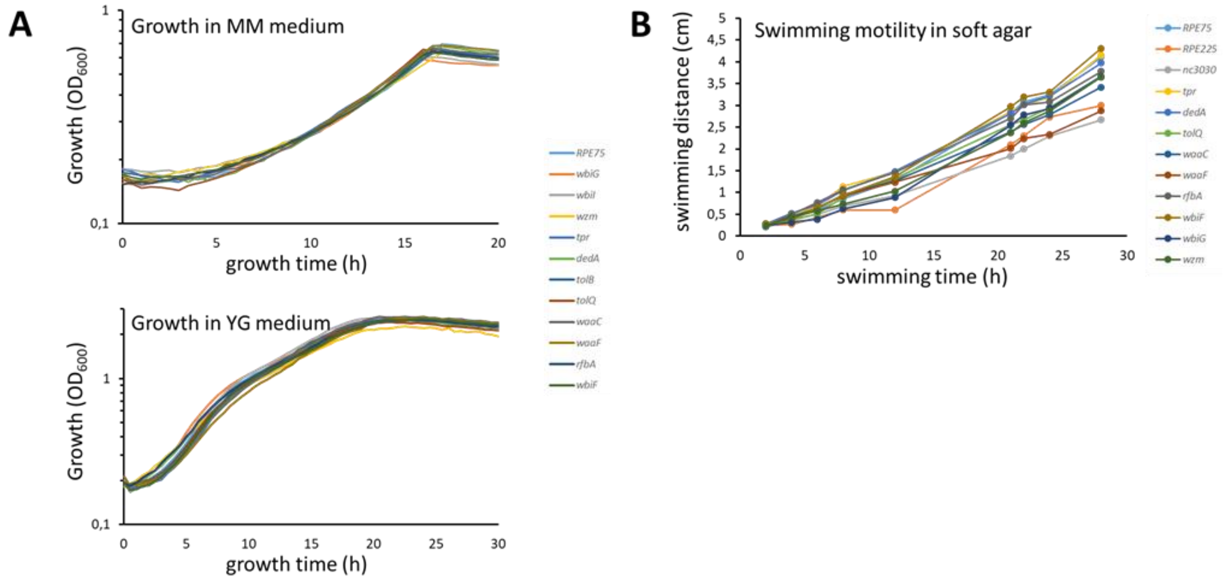


**Growth characteristics of the *C. insecticola* wild type and mutants.** (**A**) Growth curves of *C. insecticola* wild type (RPE75) and the indicated mutants in MM medium (top) and YG medium (bottom). (**B**) Swimming activity of *C. insecticola* wild type (RPE75) and the indicated mutants in YG soft agar 0,3 %. The swimming distance corresponds to the diameter of the growth halo on the plate.
